# Representations converge as brain maps diverge along the cortical hierarchy

**DOI:** 10.64898/2026.02.12.702420

**Authors:** Bogdan Petre, Martin A. Lindquist, Tor D. Wager

## Abstract

Brain maps (e.g. topographies like retinotopy, somatotopy) vary across individuals. This is thought to reflect computational differences, motivating personalized functional localization. However, artificial neural networks (ANNs) show that similar performance and internal representations can coexist with diverse circuit layouts. Consequently, we tested whether spatial diversity reflects computational diversity in the brain. Using task and resting-state fMRI data, we compared regional functional topographies and representational geometries–the within-individual dissimilarities among activity patterns that capture computable information. Across individuals (*n* = 414), representations converged in higher-order cortex despite substantial topographic diversity. Thus, different, individual-specific activity patterns encoded similar information. Topographic differences only tracked representational differences in regions under strong architectural constraints, such as highly myelinated sensory-motor cortices. We show this parallels ANNs which begin with random initial layouts but learn convergent representations, if architectures are permissive. This parallel raises a developmental question: do topographies and representations also have different developmental origins? Examining twins (*n* = 394), we found representations were sensitive to developmental environments and less heritable than topographies. Together, this shows that representational convergence occurs across idiosyncratic layouts in both artificial and biological systems, but is moderated by architectural constraints on circuit implementations. Accordingly, the relevance of localization-and representation-based paradigms of brain function depends on neural architecture.

## INTRODUCTION

Everyone’s brain is unique^1^. Among distinguishing characteristics, individuals show both idiosyncratic intrinsic network topographies^2,3^ and variation in canonical topographic maps like retinotopy^4^. These variants affect perception^5,6^ and motor function^7,8^, but topographic variation is greatest in association cortices^9–11^ thought to support higher-order cognition. This has led to the widespread belief that this variation reflects human cognitive diversity, motivating precision functional brain mapping^3,10–12^.

Nevertheless, the significance of individual variation in functional topographies may be overstated. Artificial neural networks (ANNs) produce some of our most successful brain encoding models^13–18^ without employing brain-like topographies. Instead, ANNs point to a different organizing principle: representational convergence. Rather than learning predictable circuit weights, different network instances learn predictable distinctions, or “representations”, of task contingencies and stimulus features. These internal representations converge in particular when architecture and training data are shared. As a result, training randomly initialized networks yields degenerate weight configurations that support the same functions^19–21^. If such representational convergence is a universal principle of neural systems, then a focus on topographic organization could come at a cost. It may misattribute neurodevelopmental variability in neural circuit implementations to task-relevant computational differences, obscuring representational structure that more directly supports behavior.

Here we asked whether variable topographies in the brain reflect variable representations, like in the canonical account of sensory-motor maps, or if topographies are an adaptive by-product of idiosyncratic initial conditions, like ANN weights. ANNs traditionally lack the physical and developmental constraints of real brains. This may permit representational convergence with more flexible implementations across ANNs. However, even when physical constraints are imposed, emergent ANN topographies remain idiosyncratic^22,23^. Further, brain topographies are most variable in transmodal cortex, which has the fewest physical constraints to bias circuit implementations, like myelination^9^ or highly differentiated laminae^24,25^. Indeed, topographic variation has been most informative for sensory traits^5,6^, whereas higher-order traits are often weakly predicted^3^ or better captured by spatially agnostic models after functional alignment of representational spaces (“hyperalignment”)^26–28^. However, the principles relating topographies to representations have not been directly investigated.

If the mechanisms underlying representational convergence in ANNs are shared with the brain, we would expect to see more common representations and flexible implementations in areas that have fewer architectural constraints. We tested this hypothesis by juxtaposing computational modeling with analyses of task and resting-state fMRI data from the Human Connectome Project Young Adult cohort (HCP-YA)^29^. We compared spatial topographies with representational geometries (state-dissimilarity matrices), examining how each varies with wiring costs and layer depth in ANNs, and whether they dissociate across cortex along analogous architectural gradients, including myelin density and network hierarchy. We contrasted these effects with other proposed determinants of topographic diversity, such as evolutionary recency and developmental timelines^9^. Finally, motivated by the role of initial conditions in shaping layouts of ANNs, we compared twins to test whether cortical topographies are more heritable than representations.

Our findings challenge a prevailing view of cortical organization. We found sensory-motor representations differed more among individuals than more abstract representations, which converged most in transmodal cortex, generalizing representational convergence previously established in ANNs to the brain. After controlling for representational variation, topographic similarity showed a strong dependence on neural architecture, and was greatest in thin (less differentiated, more densely intraconnected^24,25^), myelinated, and unimodal brain areas. Topographies were also more heritable and genetically determined than representations. Together these results show that topographic variance is best understood as an architectural prior strongly influencing sensory and motor function, but individuals idiosyncratically converge upon a common cognitive scaffold in architecturally unconstrained areas, analogous to representational convergence in ANNs. This reframes how we study the neural correlates of cognitive diversity, from the constructs we measure, to the models we build and the experiments we design.

## RESULTS

### ANN models reveal functional convergence despite topographic divergence

First we established the principle that topographies, representations and function can vary in distinct ways by extending recent work on topographic deep artificial neural networks (TDANNs)^22^. We trained multiple convolutional networks which differed only in initialization and in whether training included a spatial wiring penalty. In all models, units within each layer were assigned locations in a 2-dimensional spatial grid, but in TDANNs the wiring penalty encouraged nearby units to develop similar tuning while “ResNets” lacked this constraint.

Networks were trained using self-supervised learning on the ImageNet object recognition task. These models recapitulate many computational features of human brain circuits^13,22,30–32^, but remain small enough to train many independent instances. Resultant networks were paired into non-intersecting dyads within network-class (TDANN or ResNet) for similarity comparisons. Networks with idiosyncratic topographies learned similar deep representations. Unit activations of two example TDANNs to two example stimuli differed, and this dissimilarity compounded with layer depth (**Figure 1A**). These two networks nevertheless classified stimuli in similar ways, e.g. when they correctly classified *pholcidae* or incorrectly classified an image of a Rhodesian ridgeback as an Ibizan hound. For visual intuition about the mechanism of functional convergence, we treated each neural unit as an axis, and used multidimensional scaling (MDS) to illustrate the high dimensional activation space in two exemplary layers (**Figure 1B**). Similar stimuli were positioned at similar distances from one another in both networks’ activation spaces for both layers, indicating similar internal representations. This illustrates how networks could converge on similar discriminative decisions despite dissimilar topographies.

**Figure 1.**
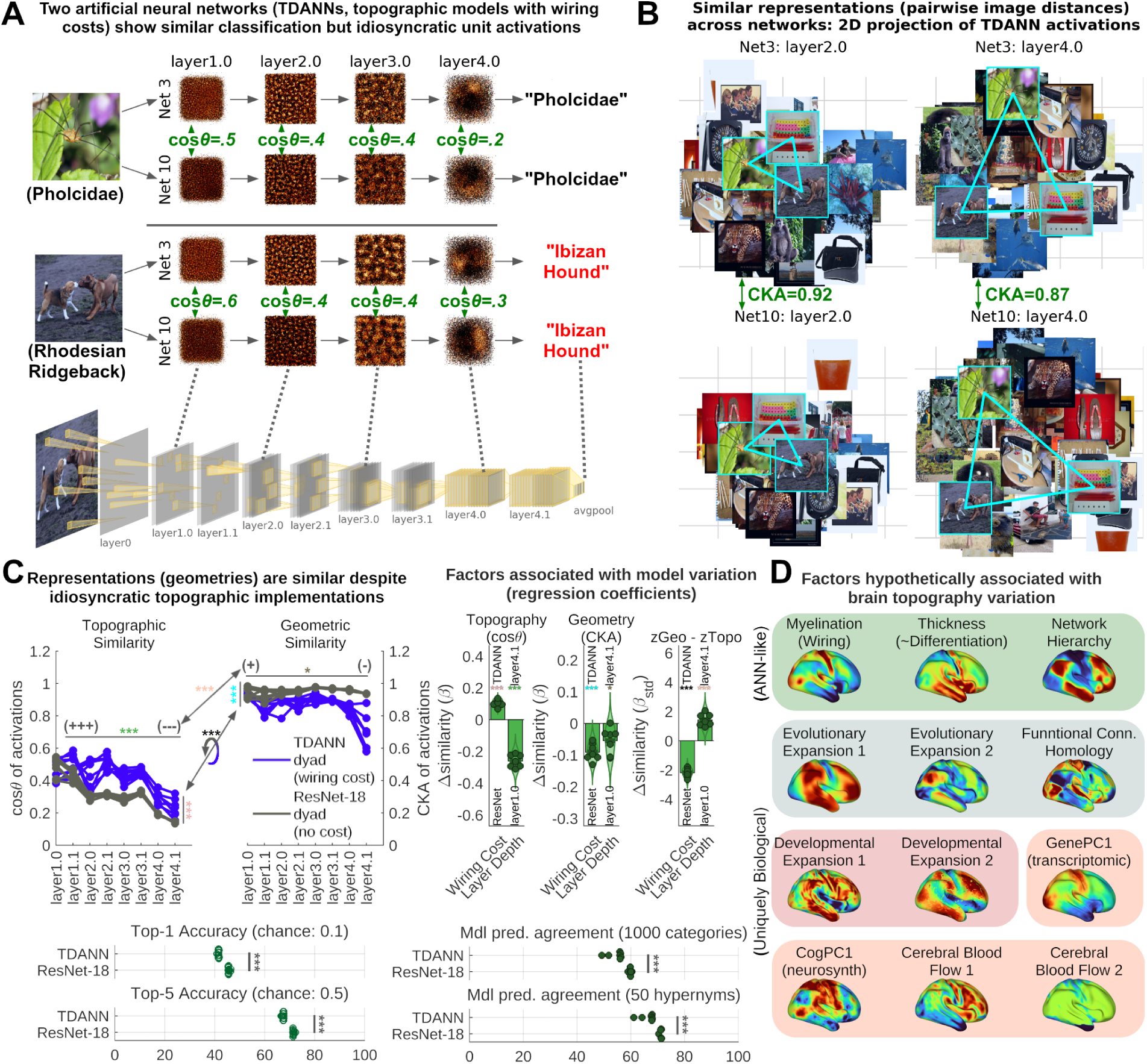
Computational experiments reveal a double dissociation of topographic and representational similarity with spatial constraints in ANNs. (A) Two TDANNs that differ only in random weight initialization seeds show idiosyncratic activations but not classifications. Select convolution layers are shown. Cos*θ* indicates spatial cosine similarity of a layer’s unit activations to an image. (B) Conservation of representational geometry is illustrated via 2-dimensional projection of the networks’ activation spaces (MDS). CKA indicates representational similarity of layers. (C) Layer depth and wiring cost (TDANNs vs. ResNets) have dissociable effects on topographic and representational similarity. TDANN dyads (*n* = 7) show more similar topographies but ResNet dyads (*n* = 7) have more similar internal representations, predictions, and greater accuracy (note: asterisks for similarities, top left, and contrasts, top right, are color coordinated). (D) Factors previously associated with topographic variation in the brain include biological analogues of layer depth (network hierarchy) and wiring costs (myelination), as well as other contending factors, used in later analyses. CKA, centered kernel alignment. *, p < 0.05, *** p < 0.001 (see **Supplemental Table 3**).

Systematic comparison of all network dyads revealed that wiring cost and layer depth had independent effects on topographic and representational similarity (**Figure 1C**, **Supplemental Table 3**). We used cosine similarity of activations to evaluate topographies. Representational similarity was evaluated using linear centered kernel alignment (CKA), a cosine-similarity-based comparison of response geometry between networks^20,33^. We found representations were very similar (CKA: 0.91 ± 0.02, mean ± se; range: [-1,1]) despite dissimilar topographies (cos*θ*: 0.36 ± 0.03, mean ± se; range: [-1,1]). Across layers, topographic similarity decayed even further, while representational similarity changed very little. The effect of wiring cost was evaluated by comparing TDANNs with ResNets, and this showed opposite effects on topographies and representations. Wiring costs facilitated topographic convergence while hampering representational convergence. This raised the question of whether topographic or representational convergence had greater functional significance.

Regardless of model initialization and obtained topographies, models performed comparably within-class (TDANN: 54.5 ± 1.1%, ResNet: 59.9 ± 0.18% mean agreement ± se), but ResNets produced greater model accuracy (TDANN: 41.5 ± 0.07%, ResNet: 45.7 ± 0.09% top-1 accuracy; top-1 accuracy *t*_13_ = 32.0, *p* = 9.3e-14) and more similar classification decisions (agreement *t*_6_ = 4.7, *p* = 3.3e-3). Notably, model decisions were even more similar to one another (model agreement) than to ground truth labels (model accuracy), indicating they were in agreement even on many misclassifications. Agreement increased further after aggregating the 1000 ImageNet category labels into 50 semantic supersets of hypernyms (TDANN: 66.4 ± 1.0%, ResNet: 71.2 ± 0.14% mean agreement ± se), meaning disagreements were often at the level of granular semantic distinctions (e.g. breed of dog rather than dog vs. house). This pattern of model agreement revealed the functional consequence of representational convergence: networks performed more similar functions where representations were most similar (ResNets), not where topographies were most similar (TDANNs).

Having established this divergence in ANNs, we next asked if similar principles hold in biological neural networks where network hierarchy and myelination may pose circuit constraints similar to layer depth and wiring costs in ANNs. Both are among the factors previously associated with interindividual topographic variation^9^ (**Figure 1D**, **Table 1**), but their relationship with representational variation remains unexplored. We turned our attention to this problem next.

**Table 1.** Neuromaps and the hypotheses they test. yo, years old.

| Neuromap | Description | Hypothesized effect | Explanation |
| --- | --- | --- | --- |
| Evolutionary |  |  |  |
| Evolutionary expansion 1 | Human vs. macaque, anatomically aligned | Phylogenetically older areas: less idiosyncratic in both topography and geometry | Older areas are more likely to have reached allelic fixation or be under greater selective pressure; newer areas would show greater genetic variation<sup>35</sup>. |
| Evolutionary expansion 2 | Human vs. macaque, functionally aligned |  |  |
| FCHomology | Functional connectivity homology vs. macaque |  |  |
| Developmental |  |  |  |
| Developmental expansion 1 | Infant vs. adult (18-24yo) areal scaling | Increased representational convergence with longer development timelines | Increased exposure to environment and susceptibility to experience dependent plasticity |
| Developmental expansion 2 | Local / global areal scaling in youth, 5-25yo |  |  |
| Architectural |  |  |  |
| Myelination | T1/T2 ratio | Greater myelination decreases representational convergence | Wiring constraint limits capacity for convergence by constraining circuit flexibility<sup>22</sup>. |
| Thickness | Cortical thickness | Representational convergence in thicker areas | Increased laminar differentiation and reduced intracortical connectivity in thinner areas<sup>24</sup> constrains circuit function. |
| Network Hierarchy | Principal gradient of connectivity | Increased representational convergence in transmodal areas | More receptive (or projective) field integration and composition increases implementation flexibility of higher order representations |
| Miscellaneous and Negative Controls |  |  |  |
| GenePC1 | Transcriptomic variation | - | - |
| CogPC1 | First principal component of cognitive terms from Neurosynth | Task dependent effects (negative control) | Task contingent effects, especially a bias favoring abstract cognitive over sensory tasks or vice versa. |
| CBF1 | Cerebral blood flow (CBF) | More consistent implementations with greater CBF | Low CBF affects the BOLD signal-to-noise ratio. |
| CBF2 |  |  |  |

### Representational geometries generalize to the brain, where they measure state discriminability

Given that representational similarity can dissociate from topographic similarity in ANNs, we first asked if the same might occur in the brain. **Figure 2** compares the same levels of analysis as **Figure 1**, topographic correspondence vs. representational correspondence, in human fMRI data. Two exemplary regions, primary visual cortex (V1) and dorsal lateral prefrontal area p9-46v, were selected in a dyad of unrelated HCP-YA participants (**Figure 2A**). Blood oxygen level dependent (BOLD) responses were evaluated across 23 task conditions spanning sensory, motor and cognitive domains, with 6 highlighted in the illustration, showing different topographic and representational characteristics in the two regions. While topographies were similar between individuals in the early sensory area V1 (spatial cos*θ* = 0.259; **Figure 2B**), they differed markedly in the higher order p9-46v (cos*θ* = 0.114; **Figure 2C**).

**Figure 2.**
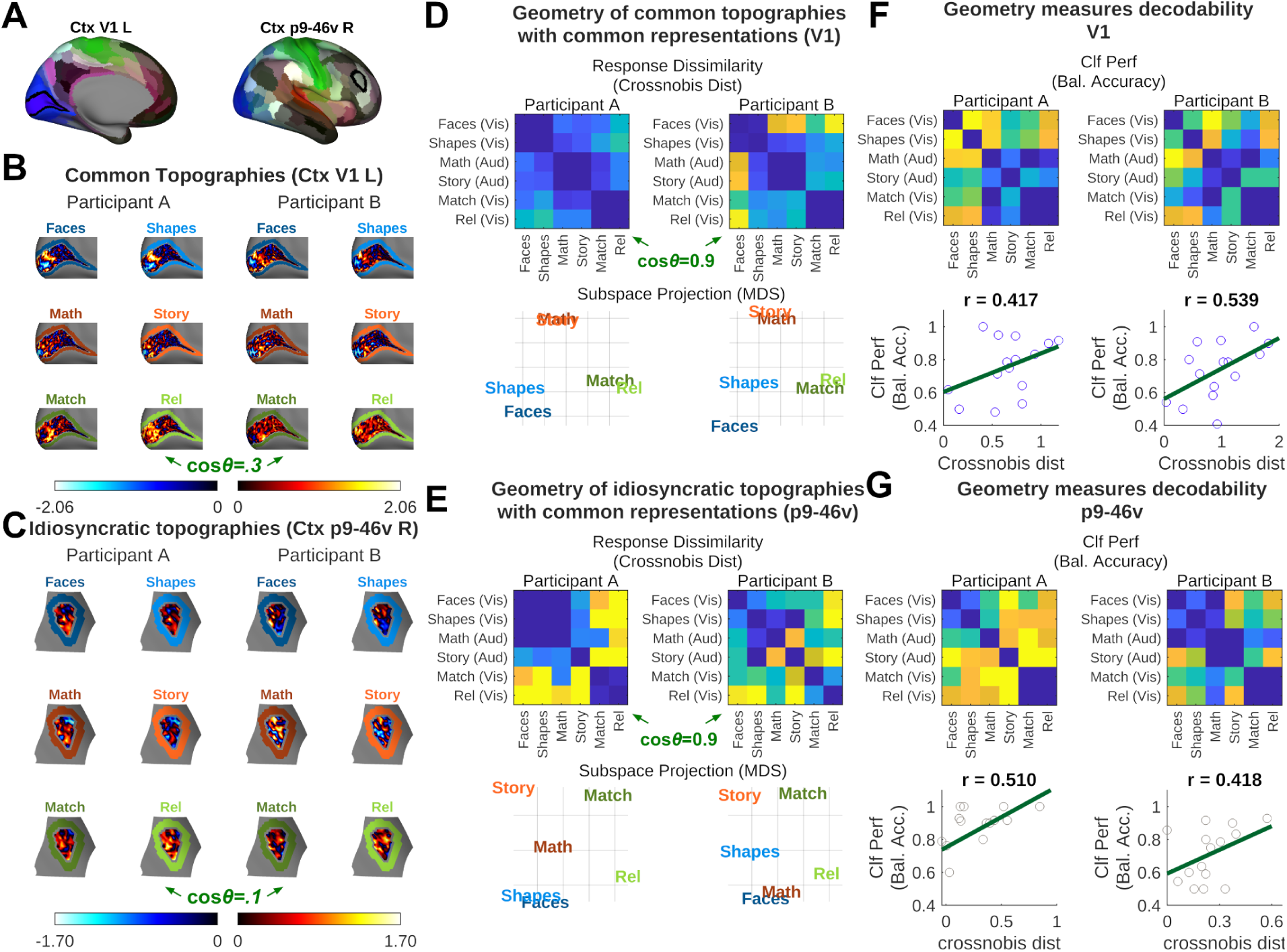
An illustrative example of common representations even when topographies are idiosyncratic. (A-C) Two exemplary regions (A: black outlines) from two participants demonstrate either similar topographies (V1) or idiosyncratic topographies (p9-46v; whitened evoked-response contrast maps). (D,E) Representational dissimilarity matrices (RDMs) capture intra-participant spatial dissimilarities. Both common and idiosyncratic topographies show common representations (compact subset of tasks shown). Cosine similarity shown on crossnobis RDMs for intuition. RDM whitening is applied later to produce WUC for inference. (F-G) Representational geometry captures task discriminability much like a brain-decoding binary nearest-centroid classifier (accuracy estimated via cross-validation across scans, weighted for condition imbalance). Each point indicates a pairwise classification vs. crossnobis distance for two task conditions. MDS, multidimensional scaling. Clf, classifier.

Representational geometry was defined in terms of multivariate distances between topographies (cross-validated Mahalanobis distances, or “crossnobis”), yielding representational dissimilarity matrices (RDMs; **Figure 2D-E**). These RDMs were very similar across participants in both regions (V1_RDM_ cos*θ =* 0.941, p9-46v_RDM_ cos*θ* = 0.915), indicating V1 had similar geometry across participants, but so did p9-46v despite its more idiosyncratic topographies. To facilitate direct comparison with our measure of representational similarity in ANNs, and account for co-dependencies across RDM rows and columns, we whitened RDMs before comparison. Whitened cosine similarity (WUC) of RDMs reduces to CKA in the deterministic limit^34^, providing a principled analog for stochastic biological data.

In the context of stochastic biological signals, the functional relevance of crossnobis RDMs lies in their relationship to state discriminability and decodability. To illustrate this and provide an intuition about our representational measures, we fit binary nearest-centroid classifiers to task evoked responses in one run and tested them on responses obtained from a second run in a 2-fold cross-validation scheme. For the 6 conditions highlighted this resulted in 15 classifiers and 15 accuracy scores, which were highly correlated with crossnobis distances (**Figure 2F-G**).

Across all 23 conditions and 207 dyads the mean correlation was consistently high for both regions (V1: *r* = 0.50 ± 0.08; p9-46v: *r* = 0.51 ± 0.09; mean ± SD), reflecting the joint reliability of classifiers and distances for these regions in this design. This means when two individuals’ brain regions have similar representational geometry the same information can be decoded from them, even when topographies differ, using individual-specific linear decoders. We characterized regional geometries using crossnobis RDMs instead of brain decoding models because they are a more robust, unbounded (classification accuracy saturates around [0,1]; *cf.* **Figure 2G**, lower left) and continuous measure of discriminability than cross-validated binary classification accuracy^36^.

### Network hierarchy dissociates topographic and representational similarity of cortical responses

We extended the analysis of topographies and representations to all brain areas to ask where they were similar or dissimilar among unrelated HCP participants (*n* = 414). Following our approach with ANNs, we paired off participants into non-intersecting dyads (*n* = 207) and evaluated within-dyad similarity. While both topographies and representational geometries showed high similarity in early sensory-motor regions, topographic similarity decayed markedly in posterior parietal cortex and dorsal lateral prefrontal cortices, whereas representational similarity remained substantially higher in these same regions (**Figures 3A-B**). Comparing standardized similarities revealed a dramatic dissociation: posterior parietal and dorsal lateral prefrontal cortices were among the least topographically similar but among the most geometrically similar regions in the cortex (**Figure 3C**).

**Figure 3.**
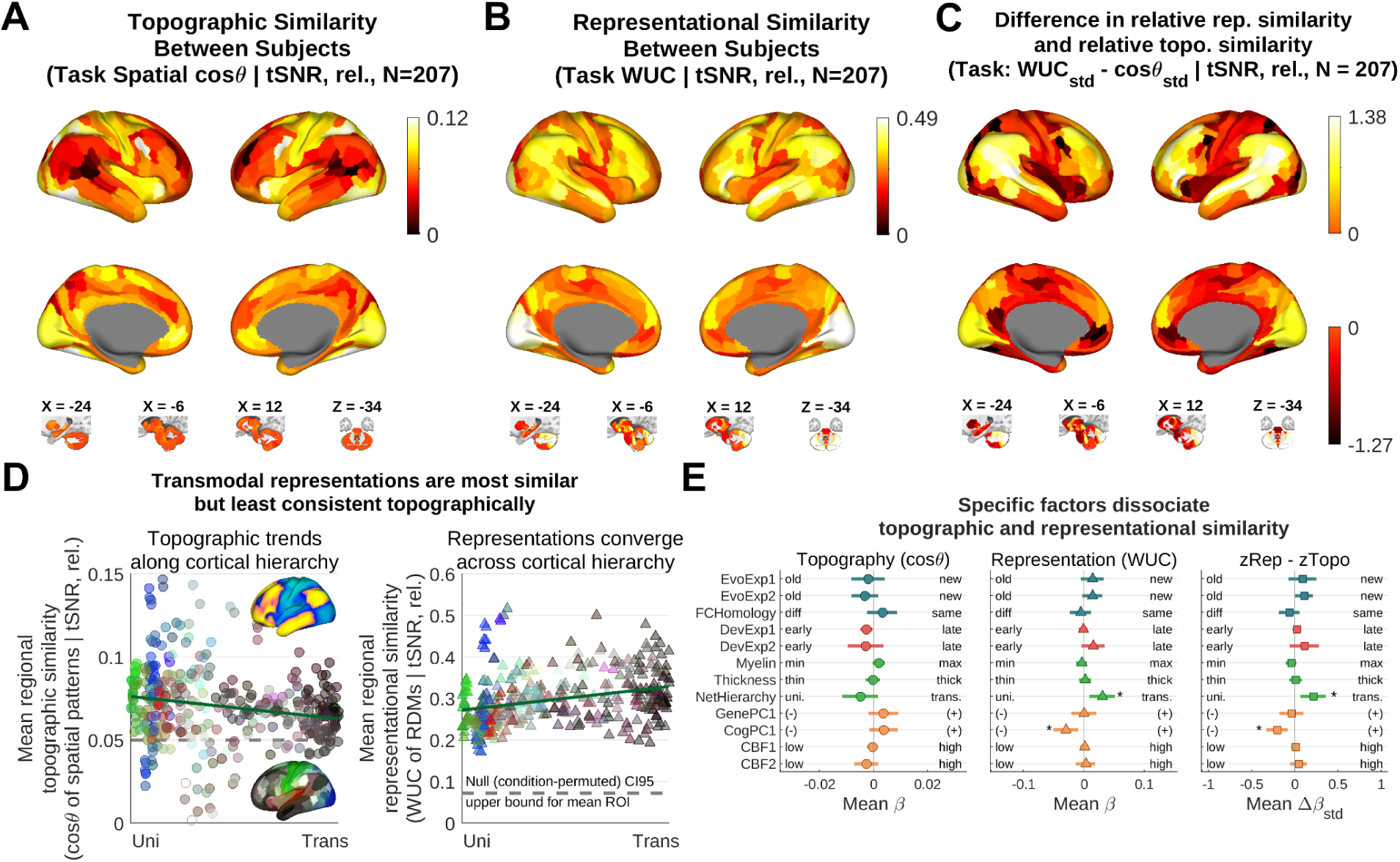
Representations are similar across participants throughout much of the cortex, especially in transmodal regions. (A,B) Topographic and representational similarity between dyads of unrelated participants across 23 different task conditions (mean spatial cos*θ* of task evoked maps and WUC of task RDMs, respectively; *n* = 207 dyads). (C) Spatial profiles of these measures differed from one another throughout the brain, indicating a dissociation between representational and topographic similarity. Metrics were standardized to enable direct comparison of spatial profiles. (D) Network hierarchy (top brain) vs. group-mean topographic similarity (left) or group-mean representational similarity (right). Topography was more consistent in unimodal areas, whereas representations were more consistent in transmodal areas. Points match the lower brain’s colors. For reference, the dashed lines indicate conservative upper bounds of the condition-permuted null similarity distributions, averaged across cortical regions (Methods). Because null similarity varies across regions, many values below this conservative reference nevertheless exceed region-specific null expectations. (E) Left, center: univariate correlations of neuromaps with topographic and representational similarity. Right: two cortical neuromaps, CogPC1 and NetHierarchy, significantly moderated the dissociation between representational and topographic similarity. Neuromaps were standardized (unit variance, zero mean across space). Association tests were limited to the cortex by neuromap coverage. Corrected for tSNR and reliability effects (**Supplemental Table 4**). Scatterplot colors: *, jackknife-adjusted spin test, FDR q=0.05. Error bars: 95% confidence interval. CI, confidence interval.

Next we inspected relationships with cortical gradients implicated in topographic diversity by prior studies^9^. Plotting topographic and representational similarity against network hierarchy (**Figure 3D**) revealed a similar dissociation as that seen in ANNs, but with a twist. TDANNs and ResNet topographies showed a dramatic decay with depth, with minimal representational depth dependence (**Figure 1C**). In the brain, topographic similarity showed slight decay with network hierarchy, whereas representational similarity increased with network hierarchy. Nevertheless, multiple prior factors have been associated with topographic diversity, including evolutionary and developmental features (**Table 1**). Each offers a plausible alternative explanation for topographic diversity, representational diversity, or both, and must be taken into account before concluding that network hierarchy offers the best explanation.

To test these competing hypotheses we introduced a jackknife adjustment to the cortical spin test (Methods: *Jackknife-adjusted spin tests*). Spin tests are association tests designed to control for spatial autocorrelation, but do not account for sampling variance across participants^37^. In Monte-Carlo simulations we found this test could be overly liberal in some cases, while a simple jackknife adjustment for sampling error resulted in a well calibrated substitute (**Supplemental Figures 3-4**). Consequently, the spatial tests we performed across the brain are more conservative than many prior studies. Additionally, we controlled for temporal signal-to-noise ratio (tSNR) and test-retest reliability in all spatial comparisons.

Systematic comparisons under this framework revealed that only network hierarchy and CogPC1 (the principal component of variation across neurosynth-derived term-maps) significantly dissociated representations from topography (**Figure 3E, Supplemental Table 4**). These factors also identified areas of high representational similarity independently (**Figure 3E**, center), but no factor was significantly associated with topographic diversity specifically. Either the more conservative inferential framework, inclusion of signal quality corrections, or choice of parcellation might explain the discrepancy with prior studies. However, the network hierarchy effects were robust regardless of parcellation (**Supplemental Figure 5E**) or of whether or not noise confound corrections were made (**Supplemental Table 4**). Together this shows that task representations converge least in unimodal areas, including higher order visual, auditory and sensory-motor regions, and converge more with greater network hierarchy, mirroring the dissociation observed in ANNs.

### Topographic similarity is only sensitive to geometric similarity where architecture constrains neural implementations

If topographies reflect underlying information content, then topographic similarity should covary with representational similarity. Therefore we used across-dyad regression to test how sensitive topographic similarity was to representational similarity. We found different sensitivity in different brain areas. For example, in V1 individuals with more similar topographies had more similar representations. Conversely, area p9-46v showed no relationship between topographies and representations despite substantial variation in representations (**Figure 4A**). Across regions there was a broad spectrum of topographic sensitivity to representational diversity (**Figure 4B**). Topographic differences were sensitive to representational differences in regions that were more myelinated, thinner, and more unimodal. Conversely, topographic differences in unmyelinated, thick, transmodal regions were not explained by representational differences (**Figure 4C**, **Supplemental Table 4**, rightmost column). GenePC1 (the principal component of transcriptional variation across the cortex) was also associated with sensitivity, consistent with its strong enrichment for myelin-related markers^38^. Together, these results show that response patterns in task-evoked fMRI primarily reflect underlying representations in regions where cortical architecture constrains how representations can be implemented.

**Figure 4.**
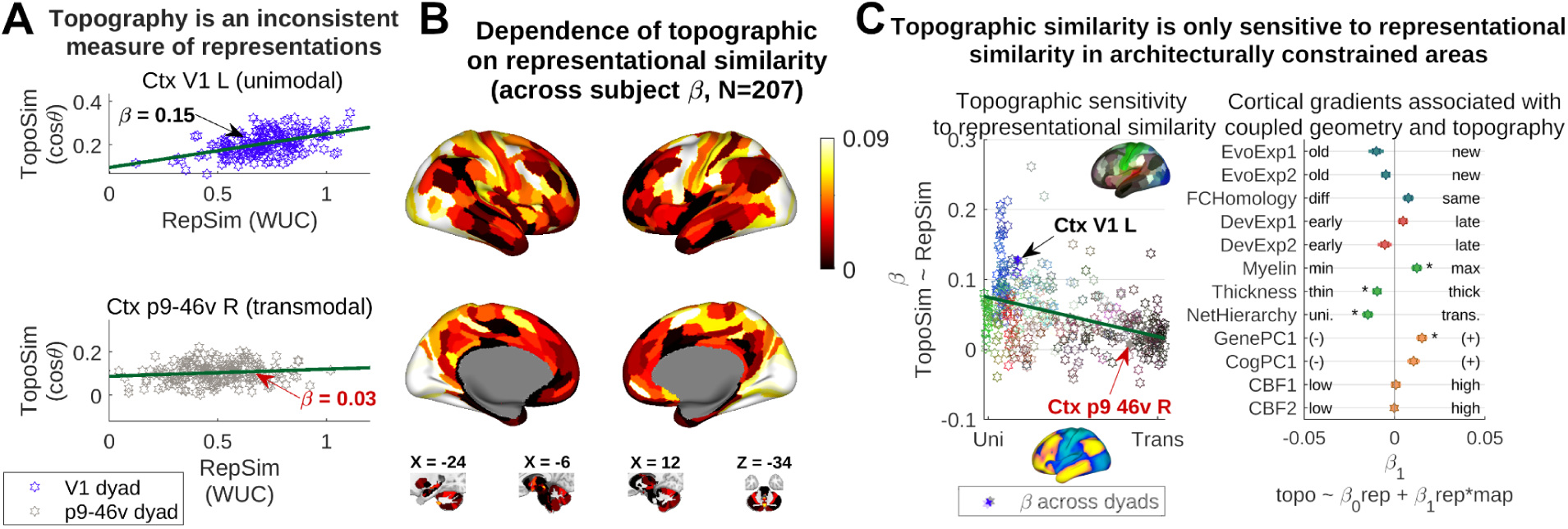
Region-by-region, topographic similarity is sensitive to representational similarity in unimodal, but not transmodal regions. (A) Two example regions illustrate how topographic similarity is sensitive to representational similarity across dyads in some areas (V1) but not others (p9-46v). Each point indicates the similarity of the region across a dyad (z-Fisher transformed). (B) Topographic similarity was sensitive to representational similarity in unimodal areas, including the somatotopic motor areas of anterior dorsal and ventral cerebellum. (C) Regional variation in sensitivity between topographic and representational similarity. Left: slope of the topography-representation relationship, like in (A), plotted against network hierarchy. Right: neuromap moderation analysis, indicating sensitivity. There was a significant association with network hierarchy such that unimodal topography was more sensitive to representational similarity than transmodal topography (jackknife-adjusted spin test, FDR *q*=0.05). Scatterplot color scheme corresponds to atlas regions (insert). Association tests were limited to the cortex by neuromap coverage. *β*_1_ adjusted for tSNR and within-participant test-retest reliability (full equation not shown, see methods). Error bars: 95% confidence interval.

### Hierarchical dissociation of representations and topographies generalizes to the cerebellum

We next asked whether the dissociation between topography and representations observed across the cortical hierarchy generalized to a distinct neural architecture: the cerebellum. Whereas the canonical cortical microcircuit is the multi-laminar column, a putative discriminative feature detector, the cerebellum is composed of numerous, fine, time-dependent inhibitory and putatively predictive circuits. Although formal gradients are not available for the cerebellum, it has areas with known unimodal topographies, as well as areas which appear transmodal based on their functional connectivity with the cortex^39^. Additionally, like the cortex but unlike many other subcortical structures, the cerebellum has reasonably good tSNR and test-retest reliability in this dataset (**Supplemental Figure 1**). This enabled a preliminary evaluation of how general the network hierarchy principle might be in determining topographic or representational convergence.

The cerebellum recapitulated the associations with network hierarchy found in the cortex. The crura at the posterior cerebellar pole are functionally connected to transmodal cortical circuits like default mode and control networks and are thus putatively transmodal, while the more anterior lobules V and VI of the superior cerebellar surface host motor representations of hand and tongue and are putatively unimodal^39^. While these structures were matched on topographic similarity, the transmodal-like crura showed substantially greater representational similarity than the unimodal-like lobules (mean crura cos*θ* = 0.059 ± 0.009, WUC = 0.530 ± 0.009; lobules V and VI mean cos*θ* = 0.060 ± 0.007, WUC = 0.351 ± 0.007; **Figure 3A-B**). Meanwhile, in the putatively unimodal lobules, topographic similarity was more sensitive to representational similarity (mean *β* = 0.077 ± [0.060, 0.093], vs. mean crura *β* = 0.042 ± [0.025, 0.056]; **Figure 4A**). Collectively these findings support the notion that network hierarchy dissociates representational geometry from topographic similarity across neural architectures.

### Architecture dissociates representational and topographic similarity of intrinsic brain networks

We tested the generalizability of our findings beyond the 23 task conditions of the HCP-YA dataset by examining representational and topographic similarity of spontaneous brain activity. We identified 25 individualized resting-state networks (RSNs) in each of our unrelated HCP participants using *a priori* group-mean network templates as a reference (Methods: Dual Regression), and analyzed individualized networks using the same analytical framework as for task evoked-responses. Under this framework topography measures the spatial layout of a network while representational geometry measures how segregated or discriminable (within-individual) the local network layouts are. We then compared these constructs within the same participant dyads as were used for task-evoked data.

As with task responses, topographic and representational similarity of RSNs showed differing spatial profiles (**Figure 5A-C**). For clarity, we illustrate representational and topographic similarity along the network hierarchy gradient (**Figure 5D**) before examining the biological factors associated with the observed inversion. Systematic comparison showed that network topographies were more similar in architecturally constrained areas (more myelinated, earlier in the hierarchy). Meanwhile, representations were consistently more similar in areas with fewer architectural constraints (less myelinated, thicker, later in the network hierarchy; **Figure 5E, Supplemental Table 5**). This inversion was statistically significant (myelination, thickness and network hierarchy all p < 0.05, FDR-corrected q < 0.05). Representations were additionally most similar in evolutionarily recent areas (EvoExp2 and FCHomology), and were associated with CogPC1 and CBF1. Of these, only myelination and network hierarchy showed significant bidirectional effects for both topographic and representational similarity, while network hierarchy showed the largest absolute regression coefficient, mirroring its dominant association in task-evoked analyses (Δ*β*_std_ > 0.49). These results align with our task-evoked findings (**Figure 3**) and identify architectural constraints as robust indicators of representational convergence with flexible implementations across participants.

**Figure 5.**
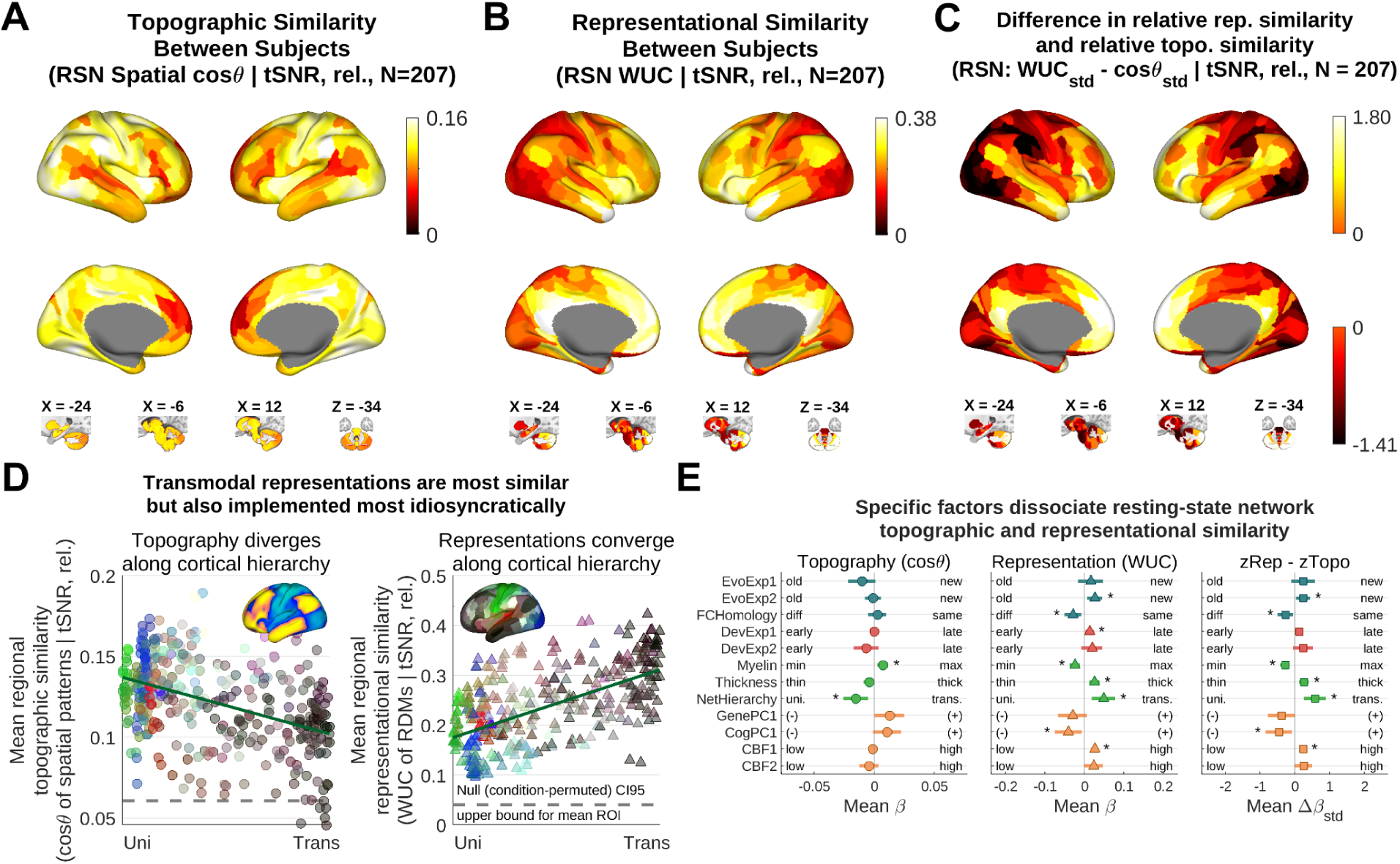
Resting state network (RSN) representational and topographic similarity diverge along architectural hierarchies, mirroring task-evoked responses (Figure 3). (A,B) RSN topographic and representational similarities vary throughout the brain in a characteristic fashion distinct from task evoked responses. Representations reflect the spatial discriminability of individualized networks within-region. Distances underlying representational geometries were computed across different scan days. (C) The spatial profile of differences between representational similarities and topographic similarities (spatially normalized within-participant to mean 0 and unit variance) was highly similar to differences between task representations and topographies (Figure 3C). (D) Network hierarchy showed the strongest dissociation of topographic and representational similarity. For reference, the dashed lines indicate conservative upper bounds of the network-permuted null similarity distributions, averaged across cortical regions (Methods). Because null similarity varies across regions, all values below this conservative reference nevertheless exceed region-specific null expectations. (E) Formal tests of the associations of neuromaps with topographic similarity, representational similarity, and their difference, revealing a tendency towards idiosyncratic implementations of common representations, reproducing task-evoked results. Association tests were limited to the cortex by neuromap coverage. Corrected for tSNR and within-participant test-retest reliability. Error bars: 95% confidence interval, see **Supplemental Table 5** for error bars smaller than icon size. *, jackknife-adjusted spin test, FDR *q*=0.05.

For completeness, we additionally inspected the sensitivity of RSN topographies to representational similarities (**Supplemental Table 5**), analogous to task-based sensitivity effects (**Figure 4**). Unlike task responses, this did not identify any dependencies with *a priori* spatial gradients for RSNs, suggesting that the observed dissociations of RSNs along cortical gradients primarily reflected population-level spatial trends rather than across-individual sensitivity of topographies to representations.

### Topographies are more heritable than representations

In ANNs matched on training data, architecture and learning rules, representational geometry converges through learning despite implementation differences due to differing network initializations^19,20^. This raises the question of whether an analogous dissociation exists between representational geometry and cortical topography in the brain. Given what is already known about the heritability of functional topographies^3,40^, we asked if representational geometry was comparatively more malleable and shaped by experience. We tested this using a twin design, comparing identical (monozygotic, MZ) and fraternal (dizygotic, DZ) twins with non-twin siblings and unrelated individuals (**Supplemental Table 1**). Genetic relatedness and shared developmental environments vary systematically across these family clusters, allowing us to dissociate genetic priors from environmental effects on cortical topographies and representational geometries. We limited ourselves to well controlled nonparametric contrasts and averages across regions. This minimized exposure to spurious nonneural confounds (Methods: *Genetic analysis*) and sample size limitations in an otherwise demanding inferential problem^41,42^.

We found topographic and representational similarity were greater among genetically related individuals, with similarity decreasing as genetic proximity declined (**Figure 6A,D**). Similarities among half-sibling task responses were an exception, but this could be explained by demographic differences (baseline gene frequencies^43^ or gene-environment interactions^44^; **Supplemental Table 1**). Similarities otherwise decayed monotonically with genetic relatedness.

**Figure 6.**
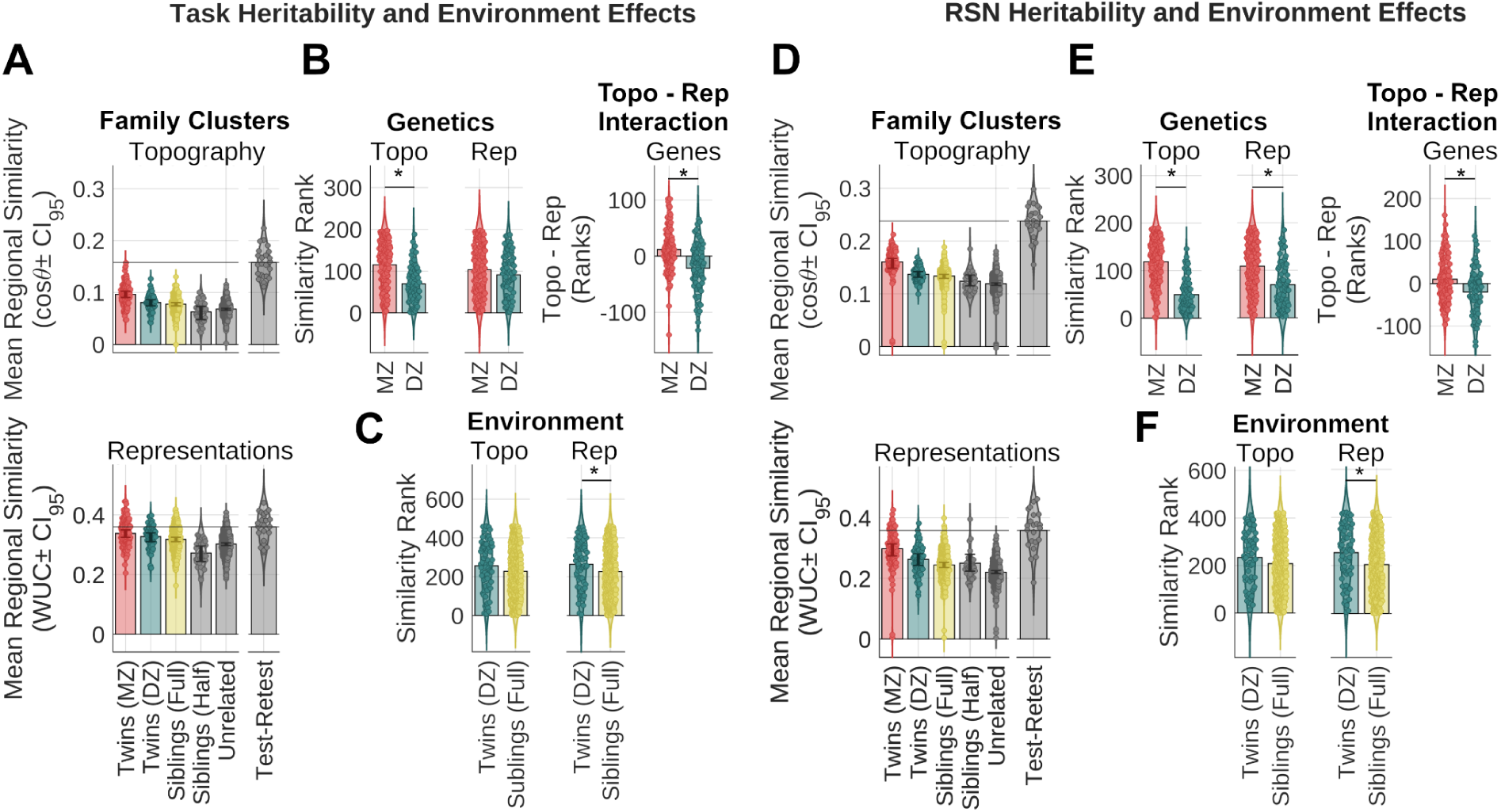
Genes exert stronger control over topography than representations, both in task responses (A-C) and resting state networks (RSNs; D-F). (A,D) Clustering individuals by familial relationships showed increasing representational and topographic similarity with genetic proximity. Test-retest reliability in individuals who returned for a second visit (*n* = 22) indicated that family relationships explain less representational than topographic variance. The horizontal line indicates mean self-similarity of retest participants, a population-average noise ceiling. This was about 2x higher for similarity of unrelated representations than unrelated topographies. (B,E) Monozygotic twins (MZ, *n* = 127 dyads) were more similar than dizygotic twins (DZ, *n* = 70 dyads), showing that both representations and topography are heritable (Mann-Whitney *U*-test, Holm-Sidak corrected for 4 comparisons, *α* = 0.05). However, heritability of topographies was significantly greater than representations in both task responses and RSNs (nonparametric interaction rank-test). (C,F) DZ twins had more similar representations than full siblings (*n* = 392 dyads), indicating a common environment effect for both task and rest (Mann-Whitney *U*-test, Holm-Sidak corrected for 4 comparisons, *α* = 0.05). Common environment effects for representations did not differ significantly from topographies for either task or RSN networks. Family clusters are color coordinated. Violin plot scatter points: similarities of non-intersecting dyads. Error bars: BCa bootstrap 95% confidence intervals (CI_95_). *, significant, *α* < 0.05.

Topographies were relatively more heritable than representations (**Figure 6B,E, Supplemental Table 6**). In both tasks and resting-state data, topographies in MZ twins were significantly more similar than DZ twins, indicating a strong genetic influence. Representational geometry also showed evidence of heritability, particularly for RSNs, but the genetic effect was significantly larger for topographies than representations in both modalities.

These differences could not be explained by differences in measurement reliability. Heritability effects were stronger for topographies than representations despite worse signal-to-noise ratio (**Supplemental Table 6)** and lower test-retest reliability (**Figure 6A,B**, gray bars).

Representations additionally showed sensitivity to developmental environments (**Figure 6C,F**, **Supplemental Table 7**). Representational geometry was more similar in DZ twins than full-siblings, who share comparable levels of genetic relatedness but differ in rearing environment. This was true for both task responses and RSNs. The common environment effect was not significant for topographies, but also did not significantly differ from representations. Thus, representations are malleable, but not uniquely so.

Together, these results indicate that cortical topography is more strongly constrained by heritable factors, and that representational geometry is also shaped by experience.

## DISCUSSION

Different research traditions variously emphasize the importance of topographic variation across individual brains or representational convergence across diverse circuits. We provide evidence of a trade off between meaningful topographic variation and representational convergence that depends on whether circuit implementations are constrained by neural architecture. Topographies are most similar in thin, unimodal and richly myelinated sensory-motor cortices, and when these topographies vary they indicate differences in how underlying information is represented. Topographies diverge most where network hierarchy is most transmodal, and in thicker, less myelinated cortex. However, rather than becoming less predictable, representations in these regions are the most consistent across individuals, consistent with their more permissive microstructural properties. We found little evidence supporting competing explanations of this implementation flexibility, like evolutionary recency or developmental timelines of brain regions. This shows that functional variation is associated with architectural constraints.

Prior evidence identifies multiple avenues by which architecture might constrain neural function. Myelinated tracts that are long distance and genetically guided^45,46^ limit the range of viable topographies^47,48^. Thinner cortex may reflect a myelination related bias^49^, but is also associated with greater laminar differentiation and reduced intracortical connectivity^24,25^ which constrains feasible computations. Meanwhile, network hierarchy is an indirect measure of polysynaptic distance from sensory and motor areas of the brain^50^. In unimodal areas long distance projections enforce topography, but as hierarchy increases, functional selectivity is increasingly determined by hierarchical composition of transformations and multimodal integration rather than spatial receptive (or projective) fields. We found that the pattern of dependence on architectural constraints broadly generalizes across the task-evoked and intrinsic resting-state networks we studied as well as across both cortex and cerebellum (which involve otherwise distinct neural architectures), consistent with a general organizing principle. Collectively, this doesn’t mean topographies are irrelevant, but it does encourage alternative approaches to functional localization when studying higher cognitive function.

Further extending this principle to artificial systems is outside the scope of this study, but our ANNs do serve an illustrative purpose. They show how compounding receptive fields permit greater implementation flexibility in deeper circuits. Additionally, by experimentally imposing wiring constraints on otherwise identical architectures, we demonstrated that architectural constraints can causally increase topographic convergence while hindering representational convergence in a controlled setting. Finally, they operationalized Marr’s levels of analysis: implementation, representation and function^51^. In our models the most similar functional output (image classification) was obtained from networks with more similar representations (ResNets), even though networks with wiring constraints (TDANNs) produced more similar implementations. This motivates an analogous framing of brain measures. In our ANNs, representations converge through learning despite idiosyncratic initial conditions. In humans, heritability analysis shows that brain topographies are more strongly constrained prior to experience, while brain representations are comparatively less constrained and more malleable.

Representations have practical significance for brain decoding which arises directly from how representational geometry is constructed. By using cross-validated Mahalanobis (crossnobis) distances and WUC, our approach preserves a meaningful zero point and compares geometric proportions rather than pattern correlations alone. Unlike traditional correlation-based representational similarity analysis^52^, which removes overall response magnitude and idiosyncratically centers each response, crossnobis distinguishes responses differing in amplitude and spatial pattern in a common reference frame (same zero point). This better reflects state decodability and more closely resembles covariance based descriptions of neural manifolds used to characterize population dynamics in electrophysiology^14,53–55^.

From this perspective, conserved representational geometry reflects conserved manifolds across individuals, while topographic similarity indicates how these manifolds are embedded in the anatomy by individual-specific population codes. These codes are heritable, and their representational significance varies systematically with architecture, rather than reflecting unconstrained initial conditions like in traditional ANNs. Accordingly, topographies warrant study and explanation in their own right^56^. However, our findings do clarify the relationship between implementation and representational levels of neural analysis, and identify a form of functional degeneracy^57,58^. This complements findings from electrophysiology showing conserved neural manifolds across animals performing similar behaviors, even when underlying circuit implementations are not directly accessible^59,60^.

Representational convergence has been shown before in both humans and ANNs. Functional alignment studies show different degrees of representational convergence across individuals^26,27^, and many ANN studies report convergent representational learning^19–21^, although its scope is contentious^61^. Our findings extend this work by showing when and under what conditions representational convergence may occur in the brain. Although substantial representational diversity persisted in our participants, representations were most similar where architectural constraints were most permissive of implementation flexibility, and in this respect more closely approached the flexibility of traditional ANNs.

Claims of representational convergence also extend to cross-architecture and brain-ANN comparisons. Under the recently proposed Platonic Representation Hypothesis all sufficiently capable neural systems may converge on a shared statistical model of reality^62^. This hypothesis partially draws support from brain-ANN alignment findings^13^, but such studies have been largely focused on visual processing^13,18,32,63,64^. Visual cortex is among the most architecturally constrained regions of cortex, with limited capacity for implementation flexibility. Thus, our findings suggest the convergence phenomenon motivating the Platonic Representation Hypothesis may be more pronounced than the existing brain-ANN literature indicates, particularly in less constrained transmodal cortex.

We hope future work will address several limitations of this study. Representations in ANNs are characterized by thousands of stimuli, and neural manifolds are characterized by many hours of recordings, whereas here we were limited to 23 different task conditions and 25 RSNs. This same dataset underlies many prior findings of individual variation in the human brain^1–4,11,28^, and is uniquely valuable for its size and family structure, but denser individual sampling^63^ is needed to more fully resolve representational geometry. At our sampling density, similar representational geometry means that these 23 task conditions (or 25 networks) are distinguished in similar ways by the same brain areas across participants, not that their full representational manifolds are equivalent.

Along with denser individual sampling, more informative task-derived behavioral measures will be important to bridge between representational and functional levels of analysis. This is straightforward in engineered systems like ANNs, where the computational objective^51^ is specified *a priori*. However, inferring the function of brain areas requires overcoming fundamental epistemic challenges associated with reverse engineering complex systems^65,66^. Nevertheless, emerging evidence suggests representational structure may provide a more direct behavioral correlate than topographies in both sensory^67^ and higher order cognitive domains^28^.

Additionally, longitudinal studies are needed to determine how representations arise and evolve^68,69^. Our data are cross-sectional, which limits inferences about representational learning to estimates of a common environment effect.

Finally, BOLD fMRI measures neural activity through a vascular filter. Ultra-high field imaging shows that hemodynamics have columnar scale precision^70^, but are also influenced by draining veins and other vascular effects^71–73^. We mitigate these factors by censoring putatively venous voxels during preprocessing and by spatial noise whitening. Further, interindividual vascular variability is unlikely to covary with the sensory-association axis, and we replicate an analogous dissociation in ANNs where no vascular filter exists. Nevertheless, corroboration from other measurement modalities will be important to fully disentangle neural and vascular contributions.

Our findings suggest how future studies of higher order brain function could adapt to target a more robust construct. Traditionally, fMRI has emphasized population-level functional maps obtained by averaging across sparse individual-level data after spatially smoothing idiosyncrasies, an approach which obscures interleaved signals^74^. Precision functional mapping addresses this through dense individual sampling. Representational analysis offers a complementary alternative particularly well suited to social, linguistic, working memory or compositional reasoning tasks like those used to engage transmodal cortex here. Like precision functional mapping it also benefits from dense individual sampling because repeated measures data are needed to estimate unbiased dissimilarities. However, experiments that include multiple functional conditions are also needed to provide a reference frame and reconstruct the underlying representational structure. Temporal information provides this structure in electrophysiology studies of neural dynamics^53,54,60^, and naturalistic viewing paradigms in brain imaging studies sometimes serve an analogous role. Response hyperalignment in particular uses time-series covariance in a reference task to define individual-specific population codes for task-evoked responses of interest^26,27^. Our results indicate that without such supplemental data the risk of misinterpretation increases in higher-order (transmodal) brain areas.

We link neural implementation flexibility to biological neural network architecture and show when and where representations converge. This links macroscale brain maps to mesoscale circuit constraints. From this view, spatial idiosyncrasies, especially in transmodal cortex, more often reflect flexible implementations than divergent computations. By bridging computational modeling, developmental neurobiology, and brain mapping, our results advance an integrated conceptual framework for the study of macroscale neural organization.

## METHODS

### Artificial neural network modeling

We used ImageNet data^75,76^ to train neural networks derived from the ResNet-18 architecture, following a previously developed protocol^22^.

Briefly, residual networks (ResNets) are a convolutional neural network architecture that achieves extraordinary depth through regularization by residual “skip” connections^77^. A suffix indicates the number of convolutional layers (here: 18), which, except for the first and last, are paired into “basic blocks” constituting our named “layers” (**Figure 1A**). All units were endowed with coordinate locations and embedded into a 2D spatial grid. In every case kernels have 3×3 unit receptive fields (layers not illustrated to scale). Depth endows the network with greater expressive capacity while convolution yields coarse spatial selectivity in early layers that is composed and integrated into spatially invariant features by later layers. Although early layers have intrinsic spatial selectivity due to the convolutional architecture, each layer also has multiple channels (features) which must be spatially interleaved (**Figure 1A**, stack widths). Channel organization dominates topographies of late layers but also affects fine topographies of earlier layers. To obtain a viable embedding topology compatible with smooth topographies, positions were selected to maximize correlation among nearby units in a pretraining task using sine grating sweeps inspired by prenatal retinal waves^78^.

Weights were randomly reinitialized after positioning, and the network was trained using contrastive self-supervised learning. This exploits the intrinsic symmetries of visual images by augmenting training data with transformations (e.g. reflection, rotation, color inversion) under which object identities are invariant^79^. The contrastive learning objective (“task” objective) is to make transformation-insensitive distinctions between images in a training batch. This differs from supervised learning because it does not require manual labeling of images, and more realistically models human infantile development. Most evaluations were performed at this stage, but to obtain model performances we froze the model trunk and used supervised learning to retrain the output (average pooling) layer.

We trained 28 network instances from 14 different seeds (0-13) with either task loss only (“ResNets”) or a composite task and spatial loss (“TDANNs”). Spatial loss encourages monotonic decay with spatial distance of the pair-wise correlation between units within-layer. This loss is weighted by a hyperparameter, *ɑ*, here set to *ɑ* = 0.25, following precedent^22^. Networks were paired into dyads within network class such that for each dyad we had an identically initialized dyad of networks using the alternative loss, allowing us to evaluate the marginal effect of spatial loss independently from the effect of initial conditions.

Convolutional neural networks have produced highly competitive encoding models of the ventral visual stream^13,30^, and ResNets are among the best^32,64^, particularly when trained using our protocols here^22,31^. Alternative topographic networks to TDANNs have recently been introduced^23,80^, but have been less thoroughly characterized and benchmarked. Additionally, the combination of pretraining and task loss objectives endow TDANN topographies and learning mechanisms with greater biological realism than alternative models. Rather than deeper ResNets, we used ResNet-18s as the base architecture for expedient training (40-80 hours/seed; Supplemental Methods: *Computational Modeling*).

Models were implemented using the pytorch framework. Model topographies and representational geometries were evaluated using post-residual activations. In both cases we flattened units across channels and either computed cosine similarity (topographies) or used the unbiased minibatch CKA estimator (representations), with a batch size of 500. To increase estimator precision minibatch CKA estimation was performed over the full ImageNet validation dataset across 10 independent passes with different image orderings, and CKA estimates were averaged across passes, following prior work^33^. All model evaluations, including classification outcomes, were performed using the independent validation set of 50,000 ImageNet images. Classification performance was evaluated against the 1000 category labels of ImageNet.

Semantic disagreement was assessed by aggregating 1000 ImageNet labels into 50 more general superclasses derived from the WordNet lexical ontology^81^. For each ImageNet label, the most specific hypernym ancestor was selected that subsumed at least 20 ImageNet labels as hyponyms, using hypernym relations only (excluding meronyms or other attributes). Because of variation in lexical precision among ImageNet labels (e.g. “street sign” vs. a specific breed of dog), a typical resulting superclass subsumed 50 labels (median; interquartile range: [27, 122]). Most labels were assigned to more specific superclasses within this set (median labels per superclass: 22; interquartile range: [9,26]).

### Participants and pairing

General inclusion and exclusion criteria for participants in the Human Connectome Project Young Adult (HCP-YA) dataset have been described elsewhere^82^. For specific inclusion/exclusion criteria specific to this study refer to the Supplemental Methods.

For topographic and geometric similarity analyses we introduced additional exclusion criteria. We required all participants to have: (1) complete functional MRI (fMRI) data and task timing information for all seven HCP tasks in both left-right (LR) and right-left (RL) acquisition directions; and (2) four resting state fMRI scans. 414 such unrelated participants were identified. We paired these participants into 207 non-intersecting dyads for topographic and geometric similarity analyses. For demographic details see **Supplemental Table 1**.

For heritability analyses we additionally included related participants (*N* = 1003). This included monozygotic twins (MZ; *n* = 254 individuals; *n* = 127 dyads, *n* = 118 genotypically verified), dizygotic twins (DZ; *n* = 140 individuals; *n* = 70 dyads, *n* = 65 genotypically verified), full siblings (FS; *n* = 775 individuals; *n* = 392 non-intersecting dyads), half siblings (HS; *n* = 54 individuals; *n* = 37 dyads) and unrelated individuals (Unr; *n* = 1003 individuals; *n* = 501,662 dyads). No triplets were present in this dataset. Dyads intersect because some individuals had multiple siblings and there were numerous possible pairings of unrelated individuals. Self reported twin zygosity was used for a minority of families lacking genotypic verification.

All experimental and informed consenting procedures were approved by the Washington University institutional review board (IRB). The Dartmouth College Committee for the Protection of Human Subjects was additionally consulted and waived further IRB review for this study’s analysis plan, which abides by the HCP’s Restricted Data Use Terms.

### Experimental design

A complete experimental design description is available elsewhere^29^. Briefly, subjects underwent two days of scanning during which they performed seven fMRI tasks twice (**Supplemental Table 2**). Each task (LR and RL acquisitions) was always acquired on the same day, with some tasks collected on the first day (predominantly working memory, gambling, motor) and others on the second (emotion, language, social, compositional reasoning). Two 15 minute resting state scans (LR and RL acquisitions) were also collected on each day. A subset of 22 unrelated participants returned for a later visit and repeated this protocol.

### Image acquisition and preprocessing

This study made use of the HCP’s minimally preprocessed data, which have undergone a standardized set of processing steps to remove artifacts, correct distortions, align different imaging modalities, and prepare the data for further analysis. The data are described in detail elsewhere^83,84^. We describe the acquisition and preprocessing characteristics most relevant for evaluating topographic and geometric similarity measures in the Supplemental Methods.

### General linear modeling of task evoked responses

We adopted previously described statistical designs of HCP tasks^29^ with slight modifications to improve representational similarity analysis (RSA). In particular, to improve voxel-wise noise estimation we used SPM’s FAST time series error models when estimating run-level contrasts. While HCP shares precomputed contrasts derived using FSL, FAST has been shown to outperform FSL’s time series error model especially with “fast” sub-second TR imaging protocols^85^, and was recommended for unbiased distance estimation (Jorn Diedrichsen, personal correspondence; see *Geometric Similarity Analyses* below). This improves vertex/voxel-level noise estimation, which benefits multivariate distance estimates. Aside from this change, our GLM models were identical to those of the HCP: hemodynamic response convolved task vectors, a 200s high-pass filter and an intercept were included in our design matrix.

### Neuromaps and brain parcellations

We *a priori* selected nine “neuromaps” aligned with the sensory-association axis that reflect developmental, evolutionary or architectural factors relevant to this study, along with three additional sensory-association aligned maps used as negative controls (**Supplemental Table 3**). All were previously published, adopted from the 81 maps included with the *Neuromaps* Python package^86^, and have been the focus in part or full of previous studies of sensory-association gradients in the brain^9,87^, motivating their use here.

We defined brain “regions” for evaluating topographic and geometric similarity using two brain atlases. The first is an in-house composite atlas (CANLab2024) that includes cortical regions from the HCP MMP 1.0 atlas^88^ and cerebellar regions from the SUIT cerebellar atlas^89^. Additional subcortical and brainstem structures are also included but largely out of the scope of this study due to the poor tSNR and test-retest reliability of deep brain structures in this dataset (**Supplemental Figure 1**). For further details on these structures refer to the CANLab2024 atlas in our “Neuroimaging Pattern Masks” github repository (Data and Code Availability). The HCP MMP 1.0 atlas was chosen because it was tailor made for this dataset and delimits functionally and structurally homogenous brain areas which might plausibly serve as functional modules. A second atlas^90^ (Gordon 333 cortical regions) was used to verify the robustness of our results with respect to parcel boundaries. Because this second atlas lacked subcortical areas, it was augmented with subcortical and cerebellar structures defined by the structural labeling scheme of the grayordinate standard^83^, which subdivides all gross structures except the brainstem by laterality.

### Topographic Similarity Analyses

We computed between-participant cosine similarity of regional spatially whitened contrast maps (Supplemental Methods: *Spatial Whitening*), and calculated a weighted average across tasks as our overall topographic similarity measure. Whitening accounts for spatial autocorrelation of noise that biases between-participant similarity measures, but more importantly matches our topographic similarity to our geometric similarity measure, which uniquely benefits from whitening (Methods: *Geometric Similarity Analyses*). Notably, spatial autocorrelation was estimated from GLM residuals, ensuring that whitening does not remove true spatial correlations in the signal^36^. Task weights were applied to prevent tasks with more conditions (working memory: 8 conditions, motor task: 5 conditions) from dominating those with fewer (2 conditions each). We chose cosine similarity because it is sensitive to both the spatial organization of responses and has a meaningful zero point (unlike spatial correlations), but is insensitive to overall response magnitude. This makes it robust to spurious (non-neural) vascular and hematological idiosyncrasies affecting BOLD contrast that could confound more formal (e.g. euclidean) between-participant distance measures.

For resting state networks (RSN) we computed cosine similarity of whitened maps of individualized networks (Methods: *Dual Regression*).

### Permuted nulls

As a scale reference for similarity magnitude, we computed condition-permuted null similarities for each participant dyad and region. For each dyad, condition order was permuted in one participant and cosine similarity was computed relative to the unpermuted participant. For task data, we used balanced oversampling to account for condition imbalance. Obtained similarities were averaged across participants within-region. We evaluated 100 permutations and the 97.5% upper bounds (across permutations) were averaged across regions and plotted in **Figures 3D** and **5D** as a conservative reference scale.

### Geometric Similarity Analyses

Geometric similarity was measured by whitened and unbiased cosine similarity (WUC) of balanced RDMs. This involved estimating unbiased Mahalanobis distances (or “crossnobis” distance; defined below) between task contrasts (or RSNs), whitening the resulting representational dissimilarity matrices (RDMs) to account for dependencies across condition-pairs (not to be confused with spatial whitening), balancing across tasks by oversampling underrepresented tasks (omitted for RSNs), and computing the cosine similarity of resulting RDMs. We detail each step below.

### Spatial whitening

Mahalanobis distance is a measure of Euclidean distance after spatial whitening. This captures decodable information more effectively than raw regression coefficient (“beta”) maps by downweighting channels that covary or contain noise^36^. Spatial covariance was estimated from GLM residuals using Ledoit-Wolf regularization^91^ (Supplemental Methods: *Spatial Whitening*).

### Unbiasedness

Mahalanobis distances are positively biased, e.g. identical contrasts will have non-zero distances proportional to their measurement error. To correct for this we partitioned task data by acquisition direction (LR or RL), computed the whitened contrast difference *a_i_* - *b_i_*in each partition, and took the product across independent partitions (*a_LR_* - *b_LR_*)^T^(*a_RL_* - *b_RL_*) for all conditions *a* and *b*. This cancels out independent (e.g. thermal) noise and yields an RDM of cross-validated Mahalanobis or “crossnobis” distances that are unbiased. Positive distances indicate that task conditions were reliably distinct, while indistinguishable conditions are symmetrically distributed around zero. We partitioned resting state data by day rather than acquisition direction to increase scan availability and ensure robustness with respect to day-to-day fluctuations in connectivity.

### RDM Whitening

Contrast-by-contrast RDMs have numerous codependencies which bias between-participant RDM comparisons if unaccounted for. This includes shared physiological noise sources (e.g., caffeine intake and its effects on vasoreactivity), shared statistical error (e.g., intercept or “baseline” estimation error), and experimental-design-induced codependencies (e.g., in fusiform face area, distances taken with respect to the emotion-faces condition will correlate with distances of the 0-back memory condition that also uses face stimuli). Even if all conditions were independent, all distances in a row of an RDM share a common comparator, introducing codependence. We empirically estimated this complex covariance structure *V* for each subject and region following a previously developed approach^34^ (Supplemental methods: *RDM Covariance Estimation*), pooled covariances across participants using a standard degrees-of-freedom (*dof*) weighing, *V* = (*dof_1_V_1_* + *dof_2_V_2_*)/(*dof_1_ + dof_2_*), and whitened RDMs in each comparison using *V^-^*^1^.

### RDM Balancing

We used balanced oversampling of underrepresented tasks to keep conditions balanced across tasks in the RDM (Supplemental Methods: *RDM Balancing*).

### Cosine Similarity

We computed cosine similarity between participants using the vectorized lower triangular elements of the balanced, whitened RDMs. As in the case of topographies, BOLD signal amplitude is spuriously idiosyncratic across participants, which modulates the magnitude of distances. Cosine similarity is insensitive to magnitude information while preserving a meaningful zero point and thus the relative proportions between distances.

### Permuted null

A representational similarity reference scale was computed in the same way as for cosine similarities of topographies (above).

### Spatial Statistics

We performed two types of cortical spatial analysis. First, we evaluated how topographies and geometries compared with cortical neuromaps in a within-dyad setting, hereon termed ‘gradient similarity analysis’ (**Figure 3, 5**). Second, we evaluated how well geometric similarity predicted topographic similarity in an across-dyad setting, and tested how this relationship varied across the cortex, hereon termed ‘sensitivity analysis’ (**Figure 4**).

Where we controlled for confounds we used temporal signal to noise ratio (tSNR) and test-retest reliability maps as covariates. tSNR was computed using Connectome Workbench’s *cifti-reduce* function and averaged across runs. Test-retest reliability was estimated as the within-participant cosine similarity of task contrasts across data partitions (e.g., LR vs. RL), weighted to account for condition imbalance across tasks, providing an overall participant-specific measure. Notably, in the participants that returned for a second visit, this within-visit measure closely approximated test-retest reliability across visits (**Supplemental Figure 1D**). Both tSNR and reliability values were averaged within dyad for use below.

For both types of analyses, neuromap data was spatially normalized (spatial mean = 0, unit variance) to ensure consistent scaling across tests. Additionally, neuromap, tSNR, and reliability values were averaged within-region for comparison. Analyses were restricted to the cortex by neuromap data availability.

### Gradient similarity analysis

We regressed each z-Fisher-transformed similarity measure (geometry or topography) on each neuromap. To control for confounds we also included a tSNR map and test-retest reliability maps in this dyad-level model. In Wilkinson notation^92^,

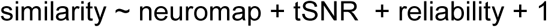

Regression coefficients were averaged across dyads. We then estimated the difference between topographic and geometric similarity and investigated how neuromaps moderated it. Here we spatially normalized similarity measures (within-dyad spatial mean = 0, unit variance) and centered tSNR and reliability confounds. Thus, reported statistics pertain to the mean cortical tSNR and reliability. In Wilkinson notation,

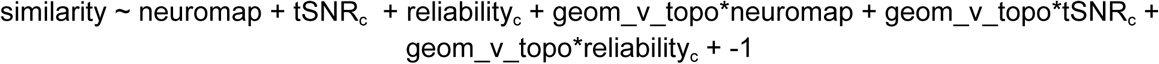

Where similarity is a normalized z-Fisher-transformed metric, geom_v_topo was coded as [0.5, -0.5] and indicated whether a similarity score indicated geometric (WUC) or topographic (cosine) similarity (respectively). Main effects and intercepts are omitted because normalization sets them both to zero by design. Inference was performed using jackknife-adjusted spin tests (below).

### Sensitivity analysis

This test evaluated how well geometry predicted topography, testing the intuition that if topographies are meaningful like retinotopy and somatotopy, then similar representations should correspond to similar topographies. We expected this correspondence to vary throughout the brain, so we tested how this association was moderated by neuromaps by fitting the following model:

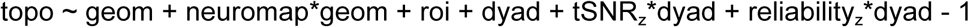

Here *topo* and *geom* indicate z-Fisher transformed topographic (cosine) or geometric (WUC) similarity measures, *roi* is a region specific intercept, *dyad* is a Helmert coded dyad contrast variable that accounts for systematic (ie brain wide, not region specific) between-dyad differences, and -1 indicates the omission of a global intercept. *tSNR* and *reliability* were spatially normalized within dyad (spatial mean = 0, unit variance). The main effect of *neuromap* does not vary within-region and is (perfectly) colinear with *roi*. The same is true for the main effects of *tSNR* and *reliability* once systematic between-participant differences have been accounted for. Consequently we only included their interaction effects. A global intercept is collinear with *roi* and was therefore also excluded. Overall the model estimates the within-region relationship between geometry and topography while controlling for overall (across regions) between-dyad differences in topographic similarity, tSNR and reliability. Inference again used jackknife-adjusted spin tests (below).

Region specific effects shown in parametric maps and scatterplots of **Figure 4** are simple univariate regressions of *cosim* on *WUC* after Fisher z-transforming both metrics. The neuromap moderator above simply indicates how these regression coefficients change across the cortex, after accounting for confounds.

### Jackknife-adjusted spin tests

A “spin” test is a permutation test that rotates neuromaps around the cortical surface to generate a null distribution that preserves spatial autocorrelation^37^. This test does not however account for sampling variance of a statistic or neuromap association. We introduced Jackknife-adjusted spin tests, which use a jackknife estimate of sampling variance to augment the null distribution’s variance under the assumption that both sampling and null distributions are Gaussian (Supplemental Methods: *Jackknife-adjusted spin test*). For our data, the resultant test was more conservative than a spin test (Supplemental Methods: *Monte Carlo Simulations*).

Confidence intervals reported for population parameters are jackknife confidence intervals.

### Dual Regression

Our dual regression approach began with resting-state network (RSN) templates from the HCP PTN1200 release. Multiple templates were available, but we made the principled choice to use the 25-network template because it most closely matched our task count (*n* = 23 conditions). Traditional dual regression multiply regresses RSN template maps on resting-state volumes to obtain individualized timeseries, which are then temporally multiply regressed on voxel/vertex timeseries to obtain individualized RSNs. For more individualized RSNs, we further incorporated weighted regression and iterated this process six times (*Supplemental Methods*: *Dual Regression Iteration Procedure and Convergence Criteria*). This approach has previously been applied to this dataset^88^, and our version primarily differs by increasing the iteration count.

We estimated the final RSNs using SPM to match the procedures used for task topographies and geometries. The individualized time-series from the final dual regression iteration were used as regressors in a GLM analysis, controlling for white matter and cerebral spinal fluid signal as well as 24 motion confounds (6 rotation and translation parameters, their quadratics, and their corresponding derivatives). Models used FAST time-series error models and 200s high-pass filters. Topographic and geometric similarity analysis proceeded identically to task-based analyses (above), except without the procedures used to address condition imbalance across tasks because RSNs had no equivalent imbalance.

### Genetic analysis

We examined the heritability of topography and geometry, which are multivariate metric traits, but measured here using BOLD fMRI contrast. Because BOLD contrast measures of metabolic reactivity are confounded by spurious (non-neural) physiological factors, all between-participant comparisons of these traits were performed after vector normalization using either cosine similarity or WUC measures. This transformation precluded the use of traditional population genetic models based on parametric analysis of variance (ANOVA)^93^.

Parametric ANOVA separates the variance of a metric trait (e.g. metabolic reactivity, our proxy for neural activity) into independent components with known theoretical contributions (e.g. a component attributable to environmental variance and one attributable to genetic differences). Our normalization step disrupted this linear additive relationship (*σ*_a+b_^2^ = *σ*_a_^2^ + *σ*_b_^2^) since *σ*_(a+b)/||a+b||_^2^ <= *σ*_a/||a||_^2^ + *σ*_b/||b||_^2^. The precise relationship cannot be recovered without the quantities ||a|| and ||b||, the net metabolic responses, which cannot be measured with BOLD fMRI alone. Instead we estimated heritability and environmental effects more indirectly.

If *a_1_,a_2_* ∼ N(*μ*,*σ_a_*^2^*I*) and *b_1_,b_2_* ∼ N(*μ*,*σ_b_*^2^*I*) are isotropic multivariate normal random variables with *μ*,*σ_a_*^2^,*σ_b_*^2^ > 0, then E[cos*θ*_a1,a2_] > E[cos*θ*_b1,b2_] entails *σ_a_*^2^ < *σ_b_*^2^. This is because projecting *x* to the surface of the unit hypersphere (i.e., *x*/||*x*||) gives rise to a von Mises-Fisher distribution whose concentration decreases as the variance increases. Thus, lower variance will produce a higher expected cosine similarity. If *μ* = 0, both are uniformly distributed on the hypersphere. For non-isotropic variance, radial anisotropy may be important, which is lost in projection. It may be possible to loosen the isotropic variance requirement in terms of the proportion of radial vs. angular variance in the unnormalized variable, but isotropy was approximated in our case by spatial whitening of topographies (±regularization) and RDM whitening of geometries regardless. The parameter *μ* is the group mean contrast or group mean geometry, so *μ* > 0 is satisfied so long as at least one task condition produces a non-zero group mean contrast which holds in our data^29^. This implies we could compare whitened and normalized BOLD derived measures within family clusters to determine whether the unobserved unnormalized traits differ in variance between two family clusters.

We tested for genetic and environmental effects by comparing family clusters matched on either environmental or genetic variance. If *σ*_G_^2^ is the genetic variance unique to DZ twins and E[cos*θ*_MZ_] > E[cos*θ*_DZ_], then *σ*_MZ_^2^ < *σ*_Dz_^2^ and *σ*_G_^2^ > 0, because MZ and DZ twins are matched on environmental variance, including uterine environment. Similarly, if *σ*_C_^2^ is the “common environment” variance due to asynchronous gestation and upbringing unique to full siblings relative to DZ twins, and E[cosθ_DZ_] > E[cosθ_FS_], then *σ*_DZ_^2^ < *σ*_FS_^2^ and *σ*_C_^2^ > 0, because full siblings and DZ twins are matched on genetic and idiosyncratic variance. We evaluated if E[cos*θ*_MZ_] > E[cos*θ*_DZ_] or E[cos*θ*_DZ_] > E[cos*θ*_FS_] using Mann-Whitney *U*-tests for genetic and common environment effects, respectively.

We compared effect sizes between topographies and geometries using an interaction test on ranks. We pooled a similarity measure (e.g., topography) across two family clusters (e.g., MZ and DZ twins) and ranked them. We then computed the difference in rank between measures (e.g., topography vs. geometry). A Mann-Whitney *U* statistic tested the null hypothesis that the between-trait rank difference was equal for the two family clusters. Rejection of this null indicated that effect sizes differed between traits.

## DATA AND CODE AVAILABILITY

All data is available from the HCP 1200 Subjects Data Release (https://www.humanconnectome.org/study/hcp-young-adult/document/1200-subjects-data-release). The code we used to train ANN models is available from the neuroAI lab: https://github.com/neuroailab/TDANN. Additional modeling code (training configurations, evaluations, etc), model weights, CIFTI files containing spatial maps from our figures and all and code needed to reproduce our brain findings from the above minimally preprocessed data are available via our own github (https://github.com/bogpetre/geometry_vs_topography/), or OSF repository (https://osf.io/xmyrn/overview), except for the CANLab2024 atlas. This atlas includes regions with a restrictive redistribution license, and can be obtained from the Neuroimaging Pattern Masks repository (https://github.com/canlab/Neuroimaging_Pattern_Masks/), which pulls constituent regions from licensed sources and rebuilds CANLab2024 locally. Alternatively, openCANLab2024 is included with the *geometry_vs_topography* repo, does not require any further processing to use, and is identical to CANLab2024 in the cortex and cerebellum, the most relevant structures for this dataset and study.

## ACKNOWLEDGEMENTS

Data were provided by the Human Connectome Project, WU-Minn Consortium (Principal Investigators: David Van Essen and Kamil Ugurbil; 1U54MH091657) funded by the 16 NIH Institutes and Centers that support the NIH Blueprint for Neuroscience Research; and by the McDonnell Center for Systems Neuroscience at Washington University.

This work was funded by NIBIB R01EB026549-03, NIMH R37MH076136-18, and Dartmouth’s Breaking the Neural Code Cluster.

We additionally thank Jorn Diedrichsen for guidance with estimating representational similarity measures, and James Haxby, Marta Čeko and Etienne Vachon-Presseau for discussion and feedback.

## AUTHOR CONTRIBUTIONS

BP: conceptualization, methodology, software, formal analysis, investigation, data curation, writing (original draft and review & editing), visualization, project administration; MAL: writing (review & editing), supervision, funding acquisition; TDW: resources, writing (review & editing), supervision, funding acquisition.

## Notes

https://www.humanconnectome.org/study/hcp-young-adult/document/1200-subjects-data-release

https://github.com/bogpetre/geometry_vs_topography/

### Competing Interest Statement

The authors have declared no competing interest.

### Summary of Updates

Aesthetic and rhetorical updates. Title has been reworded to foreground representational convergence, minor terminology updates and homogenization throughout, in particular geometric similarity was changed to representational similarity, and network depth in ANNs changed to layer depth. Contrast was increased in figure 1A, and annotations added to figure 4. Additional revisions were made to the abstract, introduction and discussion for clarity. Finally, three tables were moved from the main text to the supplement and remaining figures and table were moved from the end of the document to inline placements at their first mention in the text.

