## Supplemental Materials for "Representations converge as brain maps diverge along the cortical hierarchy"

### SUPPLEMENTAL METHODS

*Computational modeling*

We deviated from the preexisting protocol[1] only in our computational geometry, where we used one GPU per network instead of four. Batch sizes were increased from 128 to 512 images per iteration per GPU to maintain an equivalent learning rate, and CPUs per GPU (used for input preprocessing and data augmentation) were increased from 8 to 32 to keep pace. This design required approximately 40 hours per seed using 1x NVIDIA H200 GPU or 80 hours using 1x A100 GPU. All image batch sampling was done using the same fixed seed.

*Image Acquisition and Preprocessing*

Single-echo multiband (factor 8) fMRI data was acquired at 3T with 2mm isotropic voxels, 0.72s repetition time, and 33ms echo time, along with two spin-echo images with reversed phase encoding directions used for susceptibility distortion correction (SDC). This echo time is known to yield poor contrast in the subcallosal cortex, thalamus, and basal ganglia regions with shorter relaxation times[2], as quantified by temporal signal to noise ratio maps (tSNR, **Supplemental Figure 1**). Consequently, we primarily report results for cortical structures. T1w and T2w structural scans were also acquired for volumetric alignment and cortical segmentation at an isotropic 0.7mm resolution.

Functional data underwent SDC, were realigned to a single-band reference image to correct for motion (note: no further motion correction was performed during first level modeling), bias field corrected, normalized to a global mean, and resampled to a high-resolution subject specific cortical surface mesh. We treated SDC corrected LR and RL acquisitions as repeated measures to allow for unbiased distance estimation of tasks (*Geometric Similarity Analyses*, below). This assumption holds in regions with high distortion to the extent SDC is successful at the coarse scale of regional boundaries. Otherwise, distances may be over or underestimated. The mean log Jacobian determinant maps are provided as a measure of SDC throughout the brain for transparency (**Supplemental Figure 2**), and maps of test-retest reliability of response topographies (General Linear Modeling of task evoked responses, below) are provided as a secondary indirect measure of SDC success and contrast SNR (**Supplemental Figure 1**). For resting state analyses, LR and RL scans were acquired on each day, and unbiased distances were estimated across days, obviating concerns about acquisition direction related effects.

Additionally, volume to surface sampling excluded voxels with the highest signal variance, which typically corresponds to stromal voxels. This includes large draining veins that can otherwise confound fine-scale functional localization, a primary concern of this study, by producing BOLD contrast distal from the site of synaptic activity.

Cortical surfaces were downsampled and nonlinearly aligned to the Conte69 32k mesh using Freesurfer 5.2, while volumetric data was registered to the MNI152 Nonlinear 6 Asymmetric template using FNIRT for subcortical alignment. The resulting surface and volumetric spatial sampling is on average 2mm, with slight variations across the cortical mesh. At this scale, spatial precision is primarily constrained by the hemodynamic response's point spread function at 3T (~3-4mm[3]).

We used minimally smoothed (2mm full-width half maximum) CIFTI 91k formatted "MSMAll" grayordinate data. This dataset is distinguished by an additional alignment of resting state networks and T1/T2-ratio (a measure of myeloarchitecture) maps to a study specific multimodal template by smoothly ("diffeomorphically") displacing the cortical surface. Full details are available elsewhere[4]. This procedure reduces spatial differences between isomorphic, but not heteromorphic cortical topographies. To evaluate the effect of spatial alignment we also evaluated how cortical gradients dissociated representational and topographic similarity in non-MSAll aligned evoked-response data (**Supplemental Figure 6**, cf. **Figure 3**). While topographic or representational similarity of individual regions varied relative to MSMAll data, this variation did not meaningfully affect gradient tests, suggesting any gross spatial effects of multimodal alignment were orthogonal to our *a priori* gradients of interest.

*Brain Decoding*

We also fit a second GLM model to data from two participants to train brain decoders for illustrating the equivalence between representational geometries and practical task discriminability. These models were identical to our primary GLM, except instead of modeling each task condition with a single vector we split these into separate vectors for each block of stimuli or responses. This increased the number of instances for training brain decoders. Block counts per condition are detailed in **Supplemental Table 2**. Brain decoding analysis required more precise onset times and durations than our main analysis, which was obtained from E-prime files. These were missing for 12 participants who otherwise had less informative FSL-style event files used in our main analysis. Brain decoding analysis was therefore limited to 195 dyads where both participants had E-prime files.

For every pairwise combination of task conditions, we evaluated a nearest centroid classifier's ability to discriminate between a series of binary classification problems. We trained on data acquired in one run (e.g. LR) and tested on data from a different run (e.g. RL, i.e., a 2-fold cross validation scheme), and computed a balanced accuracy score to account for label imbalance by averaging the recall on each constituent condition. These balanced accuracy scores are shown in **Figure 1F-G**. Although any linear classifier could have been used, our statistical design and cross-validation scheme resulted in some conditions where only a single label instance was available for training (e.g., the 2-back tool condition of the the working memory task only offers one instance per run) and conditions with high class label imbalance between conditions. This motivated the use of a nearest centroid classifier, which is robust to small training sets and imbalance. Classifiers were trained and tested on whitened contrast maps, with one map per stimulus block.

*Spatial Whitening*

Precise estimates of the $p$ x $p$ covariance matrix of $p$ voxels/vertices can be estimated using $m$ timepoints of GLM residuals if $m >> p$, which is rarely satisfied in fMRI. Instead, we more robustly estimated spatial covariance using Ledoit-Wolf regularization[5] towards the diagonal covariance matrix. Spatial whitening was then performed by multiplying each beta map by the inverse of the estimated covariance matrix. This procedure was repeated for each participant, region and run to spatially whiten task contrast maps and RSNs.

Diagonal covariance (full regularization) is equivalent to a traditional $t$-statistic map that accounts for univariate noise by downweighting noisy channels, but does not account for information redundancy across channels. While crossnobis distance more accurately measures contrast discriminability, existing studies on interindividual differences in cortical topographies have used unwhitened contrast maps. To facilitate direct comparison with these studies, we also present results of our analyses using $t$-distances instead of crossnobis in **Supplemental Figures 7-9**. In most cases, using unbiased $t$-distances in place of crossnobis reduced our effect sizes but did not qualitatively change our conclusions.

*RDM Covariance Estimation*

The across-element RDM covariance matrix $V_s$ for participant $s$ was empirically estimated in three steps. (1) Subtraction of the mean contrast $a_{(.)}$ from each partition specific contrast $a_i$ to obtain partition-wise contrast residuals $r_i^a = a_i - a_{(.)}$ (not to be confused with time-series residuals). (2) Computing the covariance between these contrast residuals $\sigma_{a,b} = \Sigma_i(r_i^a \cdot r_i^b/p)/(n-1)$ across imaging partitions, where $a = b$ gives the variance of a contrast, and $a \neq b$ gives the covariance between contrasts, $p$ is the number of voxels/vertices, and $n$ is the number of partitions which determines the dof of $\sigma_{a,b}$. (3) Pre- and post-multiplying this covariance matrix by contrast matrices ($C\Sigma C^T$, where rows of C are contrast vectors). Along the diagonal, the variances of the distances $(a_{LR} - b_{LR})^T(a_{RL} - b_{RL})$ are $\sigma_{d(a,b)} = \sigma_a^2 + \sigma_b^2 - 2\sigma_{a,b}$. If $a$ and $b$ are spatially independent, then the variance of their distances equals the sum of the variance of the constituent measures. If they are dependent then this value is discounted proportionally. These operations generalize to the somewhat less intuitive and algebraically more complex covariances of distances along the off-diagonals. Empirical RDM whitening has been shown to increase correct model identification by 3-12%[6].

*RDM Balancing*

Finally, we addressed contrast imbalance across tasks. We included 5 motor contrasts and 8 working memory contrasts, but only had 2 contrasts from each remaining task. To avoid overrepresentation of any one task domain in between-participant similarity measures, we computed the least common multiple (LCM) of each task's condition count (LCM of 2, 5 and 8 =

40), and oversampled each task up to the LCM. Because diagonal elements of RDMs are by definition zero, oversampling produces off-diagonal null values that would positively bias RDM similarity measures. To prevent this from happening, we substituted the mean within-task distance for these null values.

*Jackknife-adjusted spin tests*

Our null hypothesis is that the observed association between a neuromap and the sample-mean spatial map contains no information about the alignment of the neuromap to the population-mean spatial map. To test this, we introduced a new test statistic,

$$z = \beta_{obs} / \sigma_{null}$$
$$\sigma_{null} = \mathrm{sqrt}(\sigma_{spin}^2 + \sigma_{sampling}^2)$$

where $\beta_{obs}$ is the ordinary least squares (OLS) sample estimate (e.g., the effect of a neuromap on geometric similarity). It measures the true population parameter $\beta$ with sampling variance $\sigma_{sampling}^2$. Meanwhile, $\sigma_{spin}^2$ is the variance of $\beta_{spin}$, the OLS estimates of $\beta$ obtained from spin permuted neuromaps. We assume that the spatial autocorrelation structure is constant across dyads, making the spin-based null distribution of $\beta_{spin}$ independent of the sampling distribution of $\beta_{obs}$. This allowed us to model the null distribution of $\beta$ as the sum of two Gaussian random variables $\beta_{spin}$ and $\beta_{obs}$. Independent variances are cumulative[7], so the variance of $\beta_{\mathrm{null}}$ is given by $\sigma_{\mathrm{null}}^2 = \sigma_{\mathrm{spin}}^2 + \sigma_{\mathrm{sampling}}^2$. If spin and sampling variance are not independent, $\sigma_{\mathrm{null}}$ above may be inflated relative to the true $\sigma_{\mathrm{null}}$, rendering the test statistic conservative.

We estimated $\sigma_{spin}^2$ with the unbiased variance estimator of $\beta_{obs}$ under spin-permuted neuromaps:

$$s_{spin}^2 = 1/(m\text{-}1) * \Sigma(\beta_{spin} - \beta_{(.)})^2$$

where $m$ = 5000 is the number of spin permutations, and $\beta_{(.)} = \Sigma\beta_{spin} / m$. When the medial wall (which is undefined) completely encompasses a parcel after map rotation, that parcel was assigned a “NaN” value and excluded from subsequent parameter estimates. Meanwhile $\sigma_{\mathrm{sampling}}^2$ was estimated using the jackknife variance estimator:

$$s_{sampling}^2 = (n\text{-}1) / n * \Sigma(\beta_{jk} - \beta_{(.)})^2$$

where $\beta_{(.)}$ is the mean $\beta$ across $n$ jackknife samples[8]. Analytic estimates of $s_{\mathrm{sampling}}^2$ give the same estimate when dyad-level estimates are available (e.g. for gradient analyses), but the jackknife estimator is more generally amenable even in situations where analytic equivalents are obscure (e.g. sensitivity analysis).

The statistic $z = \beta_{obs} / s_{\mathrm{null}}$, the unbiased estimator of $z$, has an exotic $t$-like (fat tailed) probability density function, but approaches Gaussian in the asymptotic limit. With $n$ = 207, we treat it as such for analytic tractability, at the expense of a slight underestimation of extrema likelihoods. However, we verify that this statistic remains more conservative than the traditional nonparametric spin test in all cases (one exception, **Supplemental Figure 5D**, thickness, due to a slight asymmetry in the spin distribution, but the discrepancy is minor and nonparametric spin is still significant), and use Monte Carlo simulations to verify that the false positive rate remains well controlled (Supplemental Methods: *Monte Carlo Simulations*). Thus, the *p*-values we report for spatial analyses reflect two-sided cumulative probability densities under a Gaussian, parameterized by our z-statistic.

*Monte Carlo Simulations*

We verified the calibration of the jackknife-adjusted spin test's false positive rate (FPR) using Monte-Carlo simulations.

To generate our data distribution, we bootstrap resampled unrelated participants ($n$ = 414) to form 207 new dyads per bootstrap. We simulated the spatial autocorrelation of neuromaps by sampling from spun versions of real neuromaps. For each bootstrap, we used each of the 12 spun neuromaps, and evaluated relationships between each neuromap and our bootstrapped sample. We evaluated the relationship of neuromaps with (1) task geometry, (2) the difference between task geometry and task topography and (3) the dependence of task topography on task representational geometry using the null maps and bootstrap samples. Here (1) corresponds to the main effects of gradient analysis (**Figure 3E** and **Figure 5E**, left and center), (2) corresponds to the interaction effect in gradient analysis (**Figure 3E** and **5E**, right), and (3) corresponds to the relationship between neuromaps and the sensitivity of topographic similarity to representational similarity (**Figure 4C**). Each test was repeated 1000 times for each neuromap, resulting in 12000 tests. For sensitivity analysis (3), we also ran Monte Carlo simulations where we did not spin observed maps, but instead used the true neuromaps and permuted the region labels for topographic similarity measures in bootstrapped dyads. This disrupts the relationship between topographic and representational similarity underlying the analysis.

We evaluated FPRs under three inferential frameworks. The first framework, used only for gradient analysis simulations, was a simple bias corrected and accelerated bootstrap confidence interval test, testing the null hypothesis that a regression coefficient $\beta$ = 0. This test relies on sampling error alone and is expected to produce inflated FPR due to spatial autocorrelation and motivated the development of the traditional spin test[9]. We did not evaluate bootstrapped confidence interval tests in sensitivity analysis simulations. The second framework, the traditional spin test, was tested in all cases. In this test the observed regression coefficient is compared with the distribution of regression coefficients obtained from the spun neuromaps. The *p*-value was the proportion of spun neuromaps showing a regression coefficient more extreme than the one obtained from the neuromap tested (also a spun neuromap). Note the latter neuromap is guaranteed to have a true null-relationship with our population, but may nevertheless still show a large relationship with a particular bootstrap sample if the sampling variance, which is unaccounted for, is high. This tests the likelihood that any particular sample shows a relationship with a particular null neuromap that's unusually high relative to the other null neuromaps. The third framework was a jackknife-adjusted spin test which used a parametric z-test to calculate *p*-values based on the joint spin distribution and sampling distribution. This test accounts for both autocorrelation and sampling variance. In all frameworks we included tSNR and test-retest reliability as confounds.

We evaluated our tests across the entire range of prospective FPRs by sorting obtained *p*-values and computing the empirical FPR at each nominal $\alpha$. A well calibrated test should should give an empirical FPR equal to $\alpha$. Results are discussed below and presented in **Supplemental Figures 3-4**.

*Dual Regression Iteration Procedure and Convergence Criteria*

We began by estimating an alignment map via dual regression of resting state data against 15-dimensional and 25-dimensional HCP group ICA templates. Individualized network vertices were then (Pearson) correlated with the source ICA maps, and the two obtained alignment maps were averaged. This map indicates vertex-by-vertex the similarity between individualized RSN features and the group template's RSN features. The objective is to weigh similar vertices more heavily during spatial regression since low-alignment vertices are more likely to contain idiosyncratic connectivity. To enhance contrast, we combined unsmoothed and smoothed (sigma=14mm surface and volumetric kernels) versions of these alignment maps using ((mean(weights) + weights - smoothed_weights)*((mean(weights) + weights - smoothed_weights)>0))^3, where mean(.) indicates a spatial mean. The resultant alignment was then multiplied by an areal distortion map that weighed vertices in proportion to their corresponding (barycentric) surface area. Smoothing and barycentric areal estimation were performed using Connectome Workbench.

We iterated dual regression 6 times. This was motivated by preliminary evaluation of within-participant convergence in a development sample of 10 individuals. We found that individualized ICAs from day 1 became progressively more similar to those derived from day 2 (within-participant convergence), but diverged after the 7th iteration. On the first iteration we weighed the spatial regression by a combined alignment map and areal distortion map, while subsequent iterations were weighed by the areal distortion map alone. On the first iteration this prioritizes the contribution to individualized RSN timeseries of vertices with similar resting state features as the RSN templates from which they were derived, but after the first iteration subsequent RSN templates are derived from the participant's own data, eliminating the need for an alignment map. Meanwhile, the areal distortion map accounted for heterogeneity in mesh density throughout the cortex.

The areal distortion map only affected cortical vertices, though subcortical vertices were included throughout.

### SUPPLEMENTAL RESULTS

*Monte Carlo Simulations*

Traditional spin tests performed adequately for evaluating dissociations of topographic and representational similarity along cortical gradients, but so did jackknife adjusted spin tests. For each null hypothesis test framework we evaluated the nominal probabilities (*p*-values) obtained across our 1000 tests for each of the 12 neuromaps. By plotting the incidence of *p*-values below some threshold we could compare how nominal values compared with empirical incidences. In a well calibrated test, they should be equal. Under the bootstrapped confidence interval tests (**Supplemental Figure 3**, left) the association with neuromaps is systematically positively biased, producing a grossly inflated *p*-value relative to their observed frequency of occurrence. This reflects the autocorrelation related effect that motivated the development of the traditional spin test. Comparing the sample mean with the null distribution generated from spun neuromaps (**Supplemental Figure 3**, center) corrected for this bias and resulted in a well calibrated, or possibly somewhat conservatively biased test. The jackknife-adjusted spin test showed similar performance with the traditional spin test in this context (**Supplemental Figure 3**, right), indicating that sampling variance made a negligible contribution to this test.

When testing sensitivity of topographies to representations we found only the jackknife adjusted spin test properly controlled the false positive rate. Using the same method for comparing empirical and nominal *p*-values as when evaluating gradient tests, we found that the traditional spin test grossly overestimated probabilities of extreme parameter values with spun neuromaps and bootstrapped samples (**Supplemental Figure 4**, left). The jackknife adjustment corrected this overestimation and resulted in a well calibrated, if somewhat conservative test (**Supplemental Figure 4**, center). As a secondary check we also disrupted the association within region between topographic and representational similarity by shuffling topographic similarity measures and evaluating observed sensitivities against spun neuromaps. The jackknife adjusted spin test also performed well under this assessment (**Supplemental Figure 4**, right).

| Cohort | Unr 414 | MZ | DZ | FS | HS | Unr 1003 |
|---|---|---|---|---|---|---|
| Total amount | 414 | 254 | 140 | 775 | 54 | 1003 |
| Age (years) | 28.7±3.7 | 28.3±3.4 | 28.7±3.5 | 28.7±3.7 | 28.2±3.8 | 28.7±3.7 |
| Female | 55.8% | 58.3% | 54.3% | 51.7% | 66.7% | 53.2% |
| Modal Income:<br><10,000<br>[10k,20k)<br>[20k, 30k)<br>[30k,40k)<br>[40k,50k)<br>[50k,75k)<br>[75k,100k)<br>>100,000<br>unreported | $50-75k<br>7.7%<br>6.8%<br>14.5%<br>13.8%<br>6.0%<br>19.3%<br>15.0%<br>16.7%<br>2 | $50-75k<br>4.7%<br>6.7%<br>11.4%<br>9.4%<br>9.1%<br>20.9%<br>18.5%<br>18.9%<br>- | $50-75k<br>5.7%<br>7.9%<br>10.7%<br>10.7%<br>5.0%<br>32.9%<br>11.4%<br>15%<br>1 | $50-75k<br>5.5%<br>6.5%<br>12.4%<br>11.5%<br>11.1%<br>22.1%<br>14.2%<br>16.6%<br>6 | $50-75k<br>16.7%<br>18.5%<br>16.7%<br>9.3%<br>5.6%<br>22.2%<br>5.6%<br>3.7%<br>1 | $50-75k<br>6.7%<br>7.3%<br>13.3%<br>11.5%<br>9.9%<br>21.5%<br>13.7%<br>16.2%<br>7 |
| Education (years, mean ± SD) | 14.9±1.8 | 15.0±1.9 | 15.3±1.6 | 15.1±1.7 | 13.5±2.0 | 15.0±1.8 |
| Indigenous<br>Asian<br>Black<br>Multiple<br>Unreported<br>Hispanic/Latino<br>White (nonhisp.) | 0.2%<br>7.2%<br>14.5%<br>1.9%<br>-<br>9.2%<br>66.7% | -<br>4.3%<br>8.7%<br>0.8%<br>1.2%<br>2.8%<br>82.7% | 0.7%<br>4.3%<br>8.6%<br>-<br>-<br>0.7%<br>85.7% | 0.3%<br>5.7%<br>10.3%<br>1.5%<br>0.1%<br>9.5%<br>72.4% | -<br>-<br>81.48%<br>1.9%<br>1.9%<br>-<br>13.0% | 0.2%<br>6.2%<br>13.7%<br>1.5%<br>0.2%<br>8.9%<br>69.3% |

**Supplemental Table 1** Participant demographics. MZ, monozygotic. DZ, dizygotic. FS, full siblings. HS, half siblings. Unr, unrelated.

| Task | Description | Block Duration | Trials/ Block | Blocks /run | Contrasts |
|---|---|---|---|---|---|
| Emotion | Match angry or scared faces or one of two shapes to a reference stimulus at the top of the screen | 18s<br>18s | 6<br>6 | 3<br>3 | Faces<br>Shapes |
| Gambling | Guess the value of one of several cards in order to win or lose money. Each block is biased towards gain or loss. | 28s<br>28s | 8<br>8 | 2<br>2 | Win<br>Lose |
| Language | Listen to a story and answer a question about the subject, or a vocalized arithmetic problem and pick one of two solutions on the screen | 28s<br>12s | 1<br>1 | 4<br>9 | Story<br>Math |
| Social | video clips of objects either interacting or moving randomly. Indicate if objects are interacting or not or if unsure | 23s<br>23s | 1<br>1 | 2-3<br>2-3 | Interacting<br>Random |
| Compositional reasoning | Relational condition: identify if shape or texture of top row vs. bottom row of stimuli differ along the same dimension. Control condition: pre-block cue indicates “shape” or “texture”. decide if a single stimulus in the bottom row matches top row stimuli on indicated dimension. | 16s<br>16s | 4<br>5 | 3<br>3 | Relational<br>Match |
| Motor | Tap fingers, flex toes, swirl tongue | 12.5s<br>12.5s<br>12.5s<br>12.5s<br>12.5s | 10<br>10<br>10<br>10<br>10 | 2<br>2<br>2<br>2<br>2 | Left Fingers<br>Right Fingers<br>Left Toes<br>Right Toes<br>Tongue |
| Working Memory | Each block presents only face, place, tool, or body part images.<br>0 back: pre-block clue indicates target. Participants indicate when they see it.<br>2 back: participants indicate when stimulus matches penultimate stimulus. | 27.5s<br>27.5s<br>27.5s<br>27.5s<br>27.5s<br>27.5s<br>27.5s<br>27.5s | 10<br>10<br>10<br>10<br>10<br>10<br>10<br>10 | 1<br>1<br>1<br>1<br>1<br>1<br>1<br>1 | 2 Back Faces<br>2 Back Places<br>2 Back Body<br>2 Back Tools<br>0 Back Faces<br>0 Back Places<br>0 Back Body<br>0 Back Tools |

**Supplemental Table 2** Task characteristics

| | $\beta$ | $t$ | *dof* | $p$ |
|---|---|---|---|---|
| Topographic similarity | | | | |
| Layer 1.0 > 4.1 | -0.260 | -13.5 | 16.8 | 2.0e-10 |
| TDANN > ResNet | 0.107 | 8.5 | 29.7 | 1.7e-9 |
| TDANN layer v ResNet layer | 0.065 | 1.7 | 99.7 | 0.09 |
| Geometric similarity | | | | |
| Layer 1.0 > 4.1 | -0.054 | -3.0 | 7.9 | 0.017 |
| TDANN > ResNet | -0.091 | -7.5 | 7.8 | 7.7e-5 |
| TDANN layer v ResNet layer | -0.137 | -4.6 | 19.5 | 1.7e-4 |
| Standardized geometric > standardized topographic similarity | | | | |
| Layer 1.0 - 4.1 | 1.43 | 4.24 | 219 | 3.2e-5 |
| TDANN > ResNet | -2.2 | -12.4 | 219 | 3.2e-27 |
| Accuracy: TDANN > ResNet | | | | |
| Top-1 | -4.2 | -32.0 | 13 | 9.3e-14 |
| Top-5 | -4.2 | -34.5 | 13 | 3.6e-14 |
| Agreement: TDANN > ResNet | | | | |
| 1000-categories | -5.4 | -4.7 | 6 | 3.3e-3 |
| 50 hypernyms | -4.8 | -4.5 | 6 | 4.2e-3 |

**Supplemental Table 3** Comparison of topographic, representational, and functional convergence in TDANNs and ResNet-18s. Accuracy and agreement: paired $t$-test, paired by initialization seed. Others: Mixed model, random network slope and intercepts, Satterthwaite degrees of freedom (dof).

| | Uncorrected | | | | tSNR and test-retest reliability corrected | | | |
|---|---|---|---|---|---|---|---|---|
| | Topo | Geo | zRep - zTopo | $\beta$: Topo~Rep | Topo | Geo | zRep - zTopo | $\beta$: Topo~Rep |
| EvoExp1 | -0.018 ±0.035 | 0.008 ±0.068 | **0.235* ±0.158** | -0.017 ±0.005 | -0.002 ±0.006 | 0.014 ±0.019 | 0.091 ±0.163 | -0.010 ±0.004 |
| EvoExp2 | -0.006 ±0.020 | 0.020 ±0.035 | 0.171 ±0.146 | -0.007 ±0.003 | -0.003 ±0.005 | 0.014 ±0.017 | 0.109 ±0.103 | -0.005 ±0.003 |
| FCHomology | 0.017 ±0.022 | 0.013 ±0.038 | -0.118 ±0.152 | 0.013 ±0.004 | 0.003 ±0.006 | -0.006 ±0.017 | -0.065 ±0.121 | 0.008 ±0.003 |
| DevExp1 | **0.016* ±0.012** | **0.035* ±0.022** | -0.007 ±0.067 | 0.007 ±0.003 | -0.003 ±0.002 | -0.001 ±0.007 | 0.021 ±0.047 | 0.005 ±0.003 |
| DevExp2 | 0.001 ±0.031 | 0.034 ±0.058 | 0.166 ±0.185 | -0.008 ±0.004 | -0.003 ±0.007 | 0.015 ±0.020 | 0.111 ±0.172 | -0.005 ±0.004 |
| Myelin | **0.027* ±0.013** | **0.036* ±0.028** | **-0.122* ±0.062** | **0.021* ±0.004** | 0.002 ±0.002 | -0.004 ±0.009 | -0.040 ±0.065 | **0.012* ±0.003** |
| Thickness | **-0.025* ±0.015** | **-0.045* ±0.029** | 0.044 ±0.072 | **-0.016* ±0.003** | -0.000 ±0.003 | 0.002 ±0.010 | 0.010 ±0.071 | **-0.010* ±0.003** |
| NetHierarchy | -0.030 ±0.032 | -0.017 ±0.058 | **0.253* ±0.181** | **-0.022* ±0.004** | -0.005 ±0.007 | **0.030* ±0.021** | **0.213* ±0.147** | **-0.015* ±0.003** |
| GenePC1 | 0.044 ±0.038 | 0.076 ±0.074 | -0.096 ±0.144 | **0.025* ±0.005** | 0.004 ±0.006 | -0.000 ±0.021 | -0.039 ±0.139 | **0.015* ±0.003** |
| CogPC1 | 0.029 ±0.035 | 0.040 ±0.068 | -0.124 ±0.170 | 0.015 ±0.004 | 0.004 ±0.005 | **-0.030* ±0.020** | **-0.203* ±0.123** | 0.010 ±0.003 |
| CBF1 | 0.003 ±0.010 | 0.007 ±0.021 | -0.000 ±0.048 | 0.000 ±0.003 | -0.000 ±0.002 | 0.001 ±0.007 | 0.008 ±0.051 | 0.001 ±0.003 |
| CBF2 | 0.005 ±0.026 | 0.023 ±0.051 | 0.068 ±0.126 | -0.001 ±0.002 | -0.003 ±0.004 | 0.003 ±0.016 | 0.044 ±0.099 | -0.000 ±0.002 |

**Supplemental Table 4.** Univariate coefficients, regression of task similarity metrics on neuromaps (standardized; mean: 0, standard deviation: 1), with and without confound correction. Coefficients within (but not between) columns are on the same scale. $n$ = 207. Jackknife 95% confidence intervals shown. * $p < 0.05$, FDR q = 0.05.

| | Uncorrected (Est. ± CI 95) | | | | tSNR and test-retest reliability corrected (Est. ± CI 95) | | | |
|---|---|---|---|---|---|---|---|---|
| | Topo | Geo | zRep - zTopo | $\beta$: Topo~Rep | Topo | Geo | zRep - zTopo | $\beta$: Topo~Rep |
| EvoExp1 | 0.007 ±0.041 | 0.010 ±0.034 | 0.006 ±0.629 | 0.003 ±0.008 | -0.010 ±0.011 | 0.016 ±0.032 | 0.238 ±0.340 | -0.002 ±0.011 |
| EvoExp2 | 0.022 ±0.021 | 0.023 ±0.021 | -0.042 ±0.339 | -0.000 ±0.007 | -0.001 ±0.007 | **0.027* ±0.019** | **0.236* ±0.206** | -0.004 ±0.009 |
| FCHomology | -0.009 ±0.023 | **-0.030* ±0.023** | -0.148 ±0.355 | -0.001 ±0.008 | 0.003 ±0.008 | **-0.029* ±0.022** | **-0.268* ±0.223** | 0.001 ±0.007 |
| DevExp1 | **0.030* ±0.013** | 0.007 ±0.012 | -0.259 ±0.217 | 0.005 ±0.005 | -0.000 ±0.004 | **0.013* ±0.010** | 0.116 ±0.114 | -0.001 ±0.008 |
| DevExp2 | 0.019 ±0.034 | 0.012 ±0.029 | -0.098 ±0.511 | 0.006 ±0.007 | -0.007 ±0.010 | 0.019 ±0.027 | 0.234 ±0.287 | 0.001 ±0.010 |
| Myelin | 0.015 ±0.016 | **-0.028* ±0.014** | **-0.395* ±0.259** | 0.002 ±0.008 | **0.007* ±0.005** | **-0.025* ±0.013** | **-0.281* ±0.140** | 0.003 ±0.007 |
| Thickness | **-0.027* ±0.016** | **0.029* ±0.014** | **0.539* ±0.254** | -0.010 ±0.007 | -0.004 ±0.004 | **0.025* ±0.013** | **0.259* ±0.139** | -0.006 ±0.004 |
| NetHierarchy | -0.011 ±0.035 | **0.050* ±0.032** | 0.540 ±0.552 | 0.002 ±0.010 | **-0.015* ±0.010** | **0.049* ±0.030** | **0.576* ±0.318** | 0.001 ±0.010 |
| GenePC1 | 0.042 ±0.045 | -0.033 ±0.039 | -0.723 ±0.718 | 0.007 ±0.010 | 0.013 ±0.012 | -0.030 ±0.036 | -0.387 ±0.393 | 0.003 ±0.012 |
| CogPC1 | 0.026 ±0.042 | -0.042 ±0.038 | -0.635 ±0.715 | 0.006 ±0.009 | 0.011 ±0.012 | **-0.041* ±0.034** | **-0.463* ±0.387** | 0.003 ±0.010 |
| CBF1 | 0.011 ±0.012 | **0.026* ±0.010** | 0.106 ±0.183 | -0.005 ±0.006 | -0.002 ±0.003 | **0.026* ±0.009** | **0.238 ±0.100** | -0.006 ±0.007 |
| CBF2 | 0.018 ±0.030 | 0.018 ±0.026 | -0.042 ±0.484 | 0.005 ±0.005 | -0.005 ±0.008 | 0.023 ±0.024 | 0.245 ±0.261 | 0.001 ±0.007 |

**Supplemental Table 5** Univariate coefficients, regression of RSN similarity metrics on neuromaps (standardized; mean: 0, standard deviation: 1), with and without confound correction. Coefficients within (but not between) columns are on the same scale. *N* = 207. Jackknife 95% confidence intervals. *(boldface) $p < 0.05$, FDR q = 0.05.

| Modality | Similarity | $U_{MZ > DZ}$ | *p* | AUC | SNR (MZ \| DZ) |
|---|---|---|---|---|---|
| Task | Topography | 6498 | 4.2e-7* | 0.73 | 9.0 \| 7.4 |
| | Geometry | 4980 | 0.081 | 0.56 | 13.2 \| 10.9 |
| | Topo > Geo | 6389 | 2.0e-7† | 0.72 | - |
| RSN | Topography | 6973 | 1.2e-16* | 0.86 | 8.6 \| 12.4 |
| | Geometry | 5679 | 2.2e-6* | 0.70 | 7.4 \| 6.7 |
| | Topo > Geo | 5753 | 7.3e-6† | 0.71 | - |

**Supplemental Table 6** Heritability of task topographies and geometries. * $\alpha < 0.05$, One-sided Mann-Whitney *U*-test, Holm-Šídák correction for 4 comparisons; † $p < 0.05$, non-parametric interaction test on ranks. AUC, area under the receiver operator characteristic curve in a binary forced choice test (MZ vs DZ: chance = 0.5). SNR, signal to noise ratio: mean similarity / $\Delta CI_{95}$, BCa confidence intervals.

| Modality | Similarity | $U_{DZ > FS}$ | *p* | AUC | SNR (DZ \| FS) |
|---|---|---|---|---|---|
| | Topography | 15456 | 0.046 | 0.58 | 7.4 \| 14.8 |
| Task | Geometry | 15963 | 0.015* | 0.58 | 10.9 \| 20.7 |
| | Topo > Geo | 13956 | 0.409 | 0.51 | - |
| | Topography | 12871 | 0.058 | 0.56 | 12.4 \| 17.8 |
| RSN | Geometry | 14150 | 1.4e-3* | 0.62 | 6.7 \| 13.6 |
| | Topo > Geo | 12690 | 0.085 | 0.55 | - |

**Supplemental Table 7** Effect of common environment on task topographies and geometries. * $\alpha < 0.05$, One-sided Mann-Whitney *U*-test, Holm-Šídák correction for 4 comparisons. Both non-parametric interaction tests on ranks (Topo > Geo) are non-significant. FS, full-sibling. AUC, area under the receiver operator characteristic curve in a binary forced choice test (DZ vs FS: chance = 0.5). SNR, signal to noise ratio: mean similarity / $\Delta CI_{95}$, BCa confidence intervals.

### SUPPLEMENTAL RESULTS

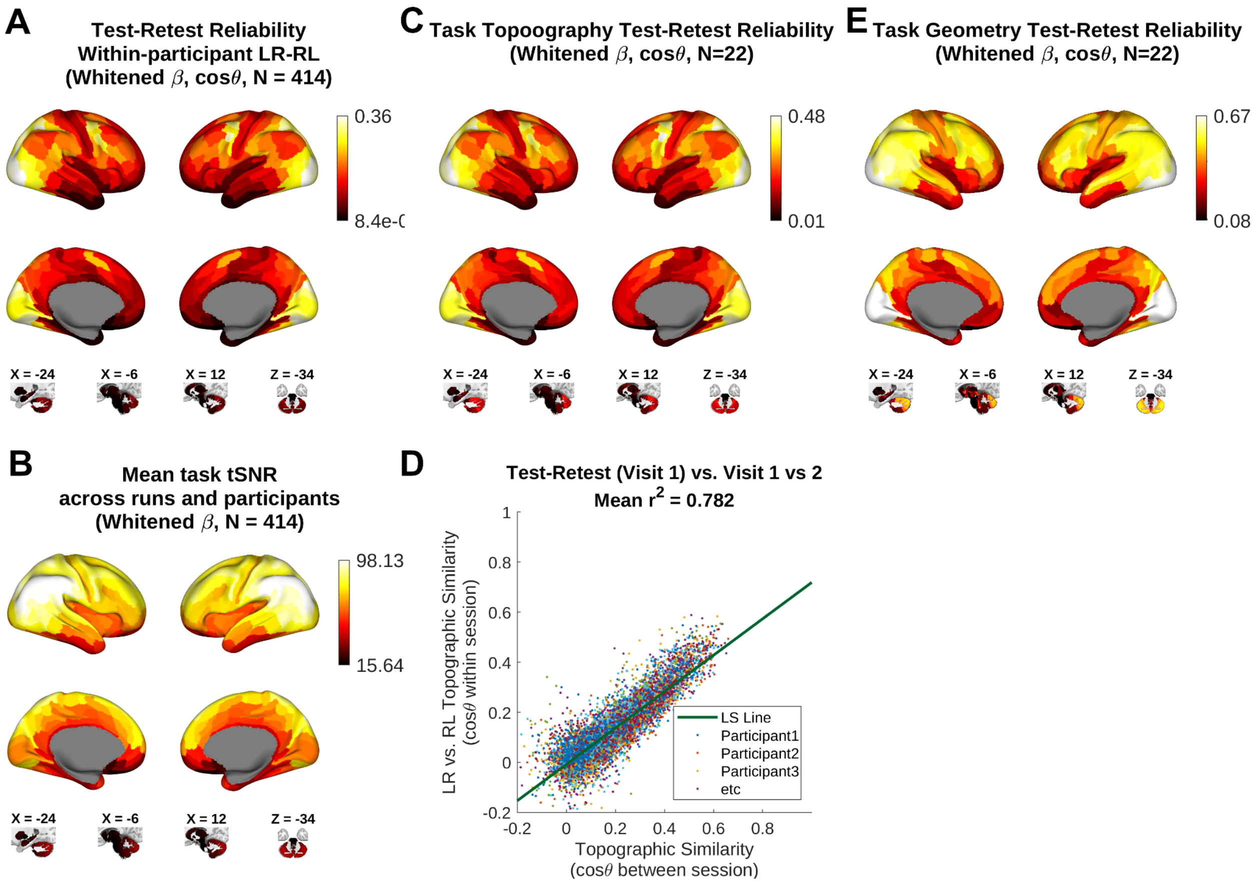


**Supplemental Figure 1** Task reliabilities. (A) Within-session test-retest reliabilities for topographic similarity of evoked responses and (B) mean tSNR across task runs serve as confound regressors in spatial analysis of neuromap associations. (C) More informative test-retest reliability was obtained across sessions for a subset of participants who returned to repeat the study (*n* = 22). (D) Within these participants between-session and between-run test-retest reliabilities were highly correlated (between-session reliability estimated at visit 1). (E) Geometry also showed good test-retest reliability throughout much of the brain between visit 1 and followup.  Test-retest reliability between visit 1 and followup serves as a noise ceiling for across participant similarities. LS, least squares.a

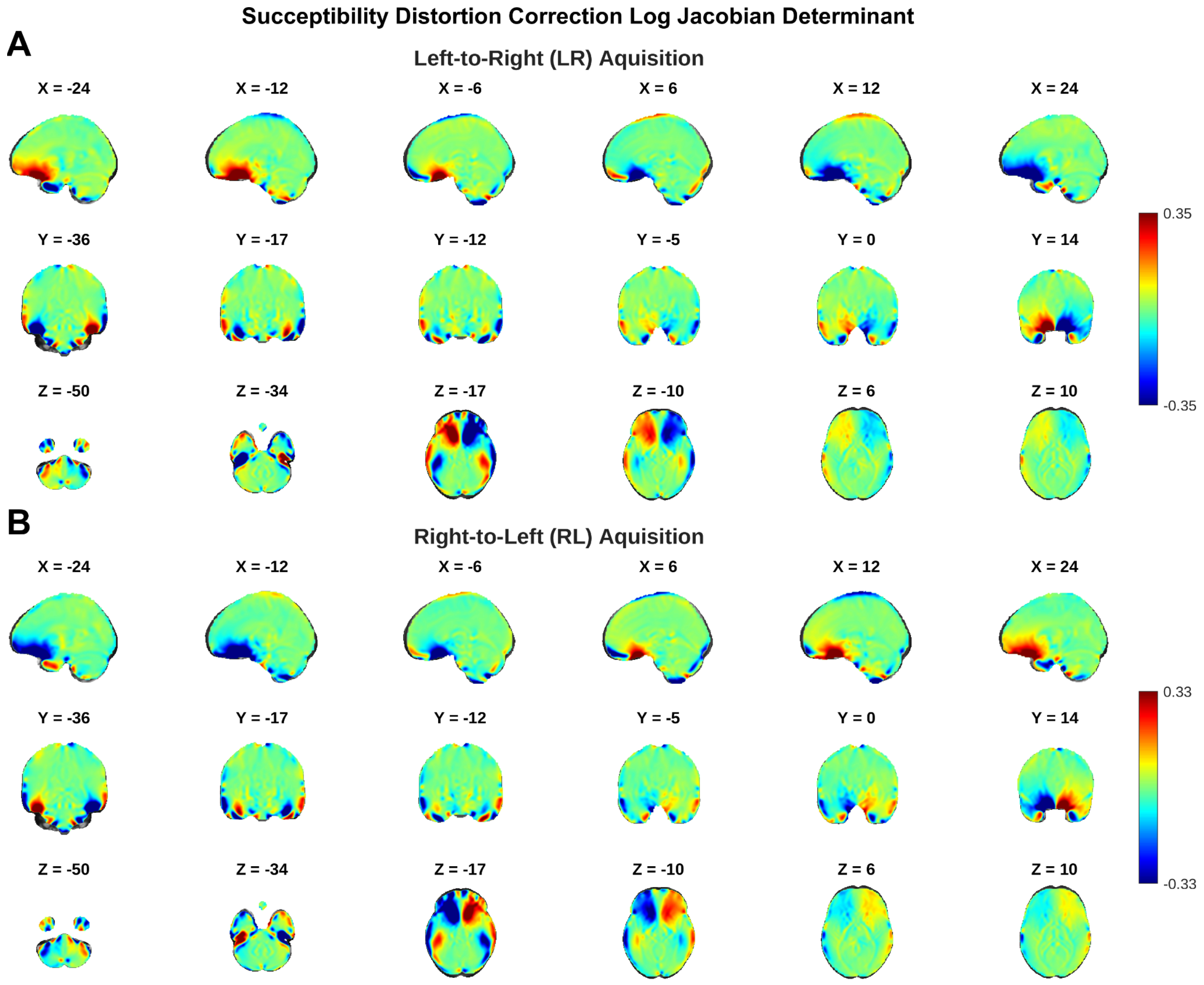


**Supplemental Figure 2** Mean log Jacobian determinants of SDCs indicates gradient nonlinearities primarily affect the orbitofrontal cortex, while the rest of the brain shows minimal distortion. This justifies the treatment of task data acquired in different directions as repeated measures in cross-validated distance estimation, because gradient direction related bias is only likely in a handful of orbitofrontal regions. MNI152NLin6Asym aligned Jacobian determinants are (natural) log transformed and summed across tasks and participants. Positive values indicate mean volume contraction, negative indicate expansion.

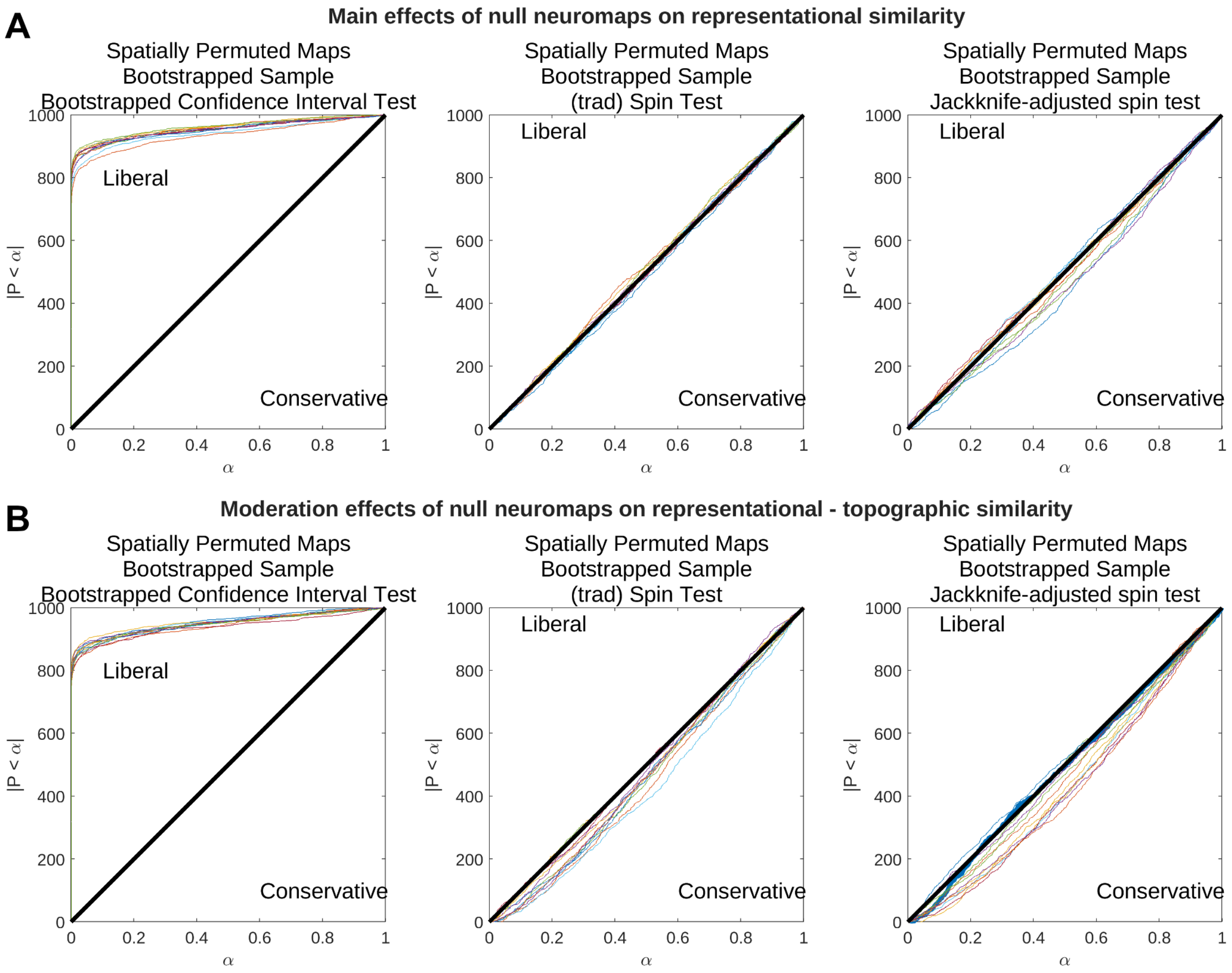


**Supplemental Figure 3** Observed (y-axis) vs. expected (x-axis) false positive rate of gradient test main effects (top) and interaction effects (bottom) for randomly rotated neuromaps drawn from the 12 neuromaps of interest (**Figure 1D**). Each colored line indicates a neuromap. 1000 bootstrapped samples (N=207) were evaluated, each tested against all 12 spun neuromaps. $|P < \alpha|$ indicates cardinality, or number of observed instances with $p < \alpha$. *t*-tests evaluate if the estimated regression coefficient is different from zero (left column), and are overly liberal, demonstrating the motivation behind the development of the traditional spin test. Traditional spin tests evaluate if the estimated coefficient is greater than that observed from a randomly rotated neuromap (center). The jackknife-adjusted spin test accounts for sampling variance in the observed coefficient (right). It tests if the population from which the sample is drawn has a relationship with the neuromap greater than that expected from autocorrelation alone. Both traditional and jackknife-adjusted spin tests show good control on the false positive rate, indicating low sampling variance for parameters in these models. Cf. **Figure 3E,5E**,**6E** **Supplemental Figure 7E,9E**.

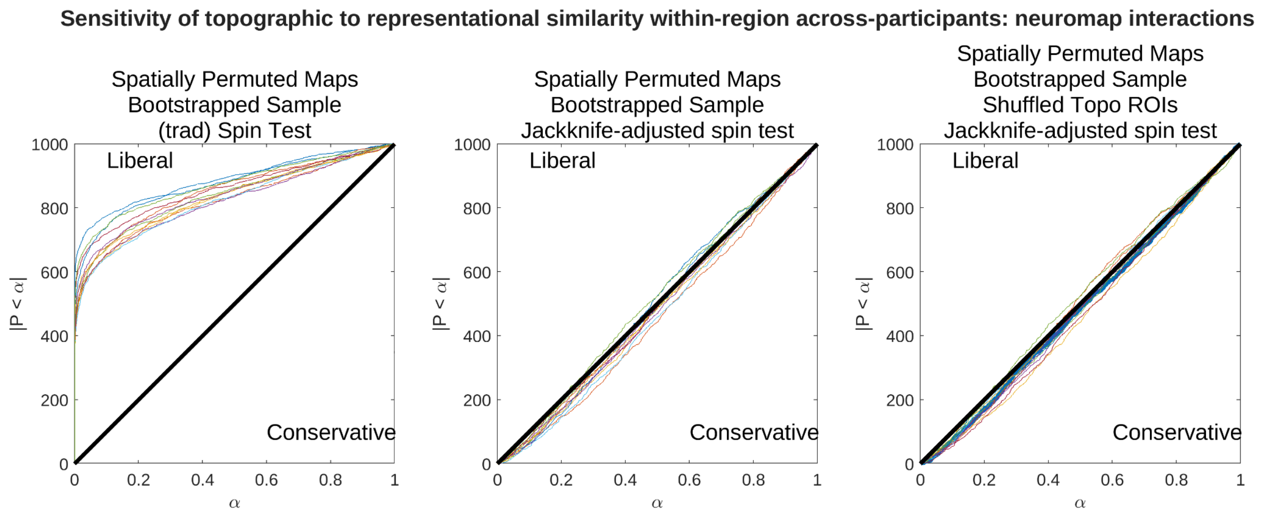


**Supplemental Figure 4** Observed (y-axis) vs. expected (x-axis) false positive rate of sensitivity analysis effects for randomly rotated neuromaps drawn from the 12 neuromaps of interest (**Figure 1D**). 1000 bootstrapped samples ($n$ = 207) were evaluated, each tested against all 12 spun neuromaps (null moderator; left, center) or with region labels shuffled for topographic similarity measures (but not geometric measures; null main-effect) and tested against the true neuromap (right). $|P < \alpha|$ indicates cardinality, or number of observed instances with $p < \alpha$. This test is performed within a region across dyads and then evaluated across regions, a test which produces significant sampling variance. The traditional spin test shows a grossly inflated false positive rate (left), but accounting for the sampling variance leads to good control of the false positive rate under the jackknife-adjusted spin test. This test continues to show good control for the false positive rate when neuromaps are held constant and region labels are instead shuffled, disrupting the relationship between representations and topography (right). cf. **Figure 4C, Supplemental Figure 8C**.

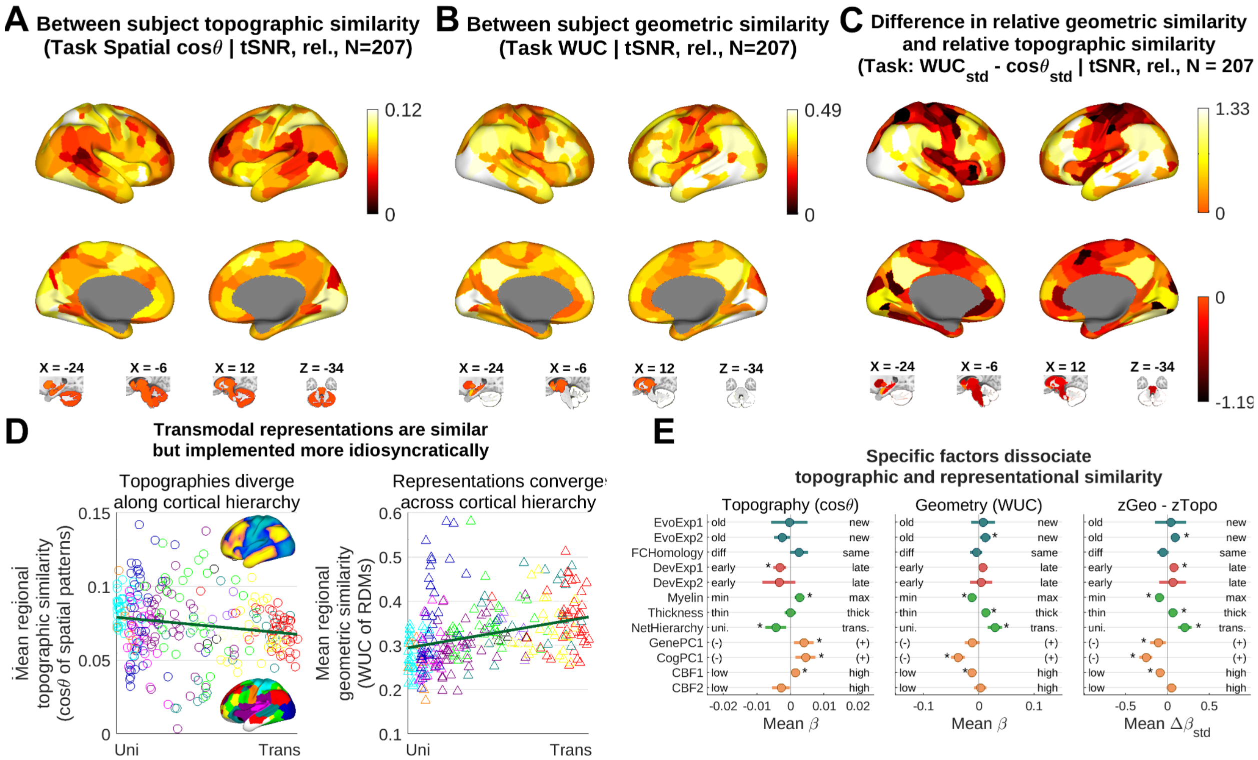


**Supplemental Figure 5** Spatial gradient effects are robust with respect to parcellation schemes. Replication of Figure 3 using the 333 parcel Gordon cortical atlas + 23 grayordinate subcortical structural labels. Topographic and representational similarity are doubly dissociated across multiple factors, including myelination and network hierarchy. Refer to Figure 3 for a description of each panel. Icon colors in (E) correspond to parcels in the Gordon Atlas (D, lower insert). Corrected for tSNR and test-retest reliability. *, $p$ < 0.05, FDR q = 0.05 jackknife-adjusted spin test. Error bars: 95% confidence interval.

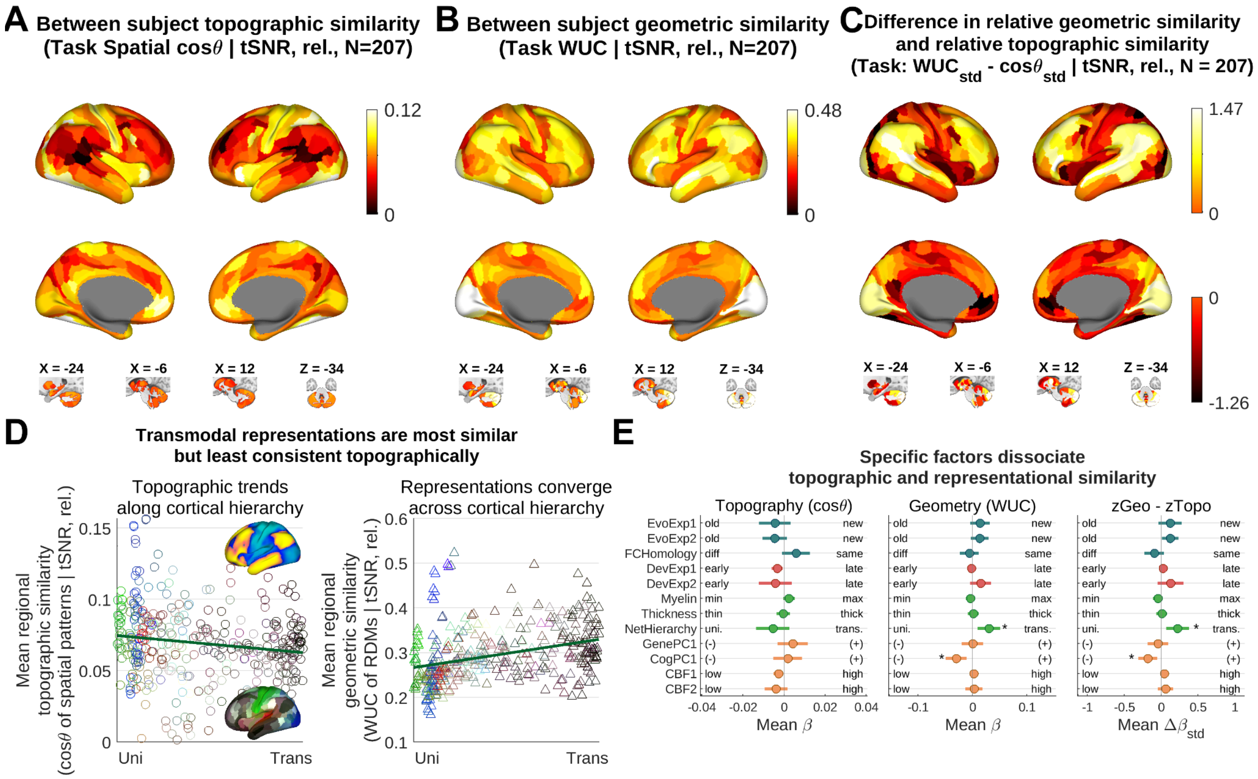


**Supplemental Figure 6** Non-MSMAll data: Dissociation of geometric and topographic similarity along cortical gradients in anatomically aligned task-evoked response data (cf. **Figure 3**). (A-C) Use of anatomical alignment alone, without “MSMAll” multimodal alignment, altered topographic and geometric similarity measures in select brain areas (e.g. V1 topographic similarity, cf. **Figure 3A-C**). (D-E) These changes did not affect topographic and representational similarity measures dissociated along *a priori* cortical gradients (cf. **Figure 3D,E**), suggesting that multimodal alignment has an effect on similarity measures that is orthogonal to these gradients and does not systematically bias our conclusions. Corrected for tSNR and test-retest reliability. * $p < 0.05$, FPR q = 0.05 jackknife-adjusted spin test. Error bars: 95% confidence interval.

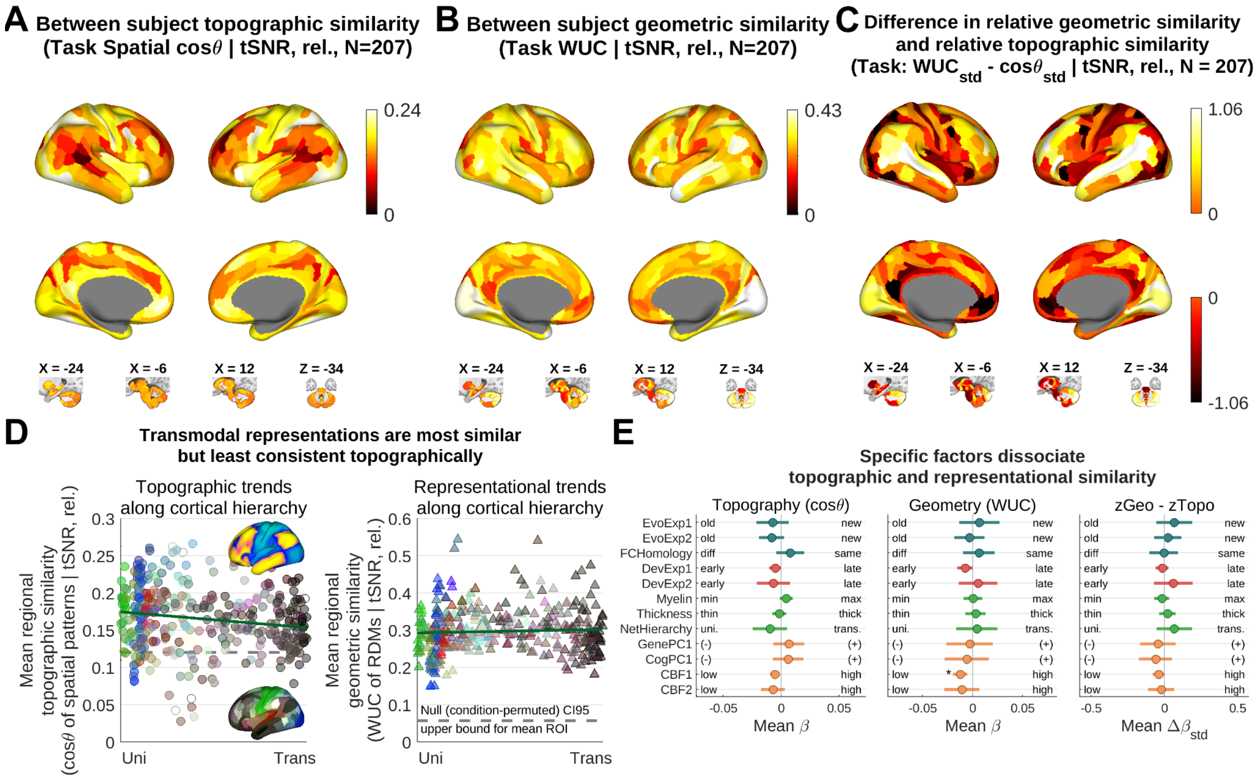


**Supplemental Figure 7** Effect of neuromaps on the difference between topographic similarity of t-statistics and WUC of *t*-distances (cf. **Figure 3)**. (A) Foregoing spatial whitening increased spatial similarities within dyads and qualitatively altered the contours of the most dissimilar regions in the parietal cortex subtly. (B) Substituting *t*-distances for crossnobis distance in RDM estimation ‘flattened’ similarity estimates across parietal cortex and dlPFC. (C) *t*-statistic based measures reduce the difference between topographic and geometric similarity in association cortex while increasing differences around visual area MT. (D) *t*-distances reduce the variance in similarities relative to whitened contrasts. (E) No neuromaps show significant associations with similarity measures using *t*-distances. This is the only instance in which we did not reproduce findings found using whitened topographies and crossnobis RDMs. However, the trends are similar, especially for the contrast of similarity and geometry (right). Associations are corrected for tSNR and test-retest reliability (also estimated using *t*-statistics in this case). *, $p < 0.05$, FDR q = 0.05 jackknife-adjusted spin test. Error bars: 95% confidence interval.

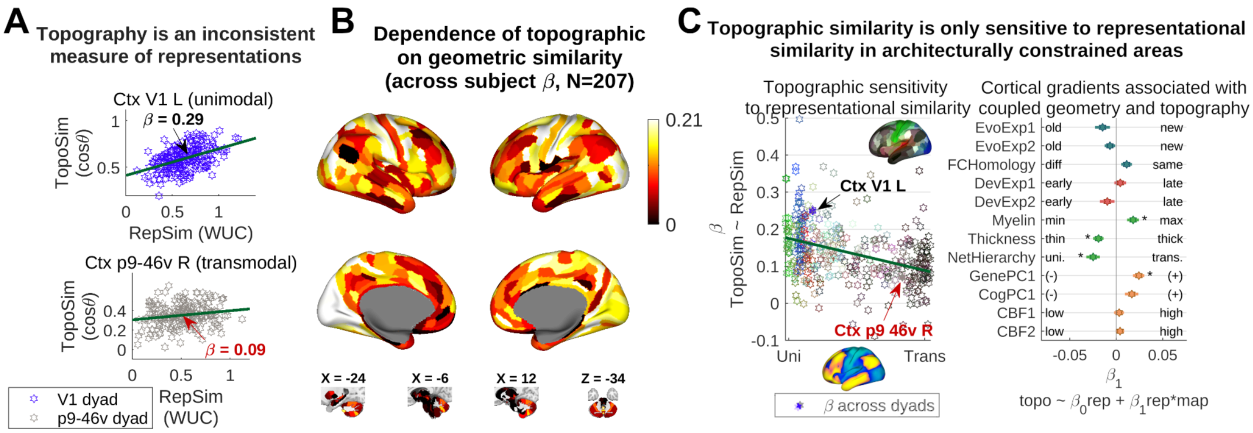


**Supplemental Figure 8** Topographic similarity is predicted by geometric similarity in sensory but not association cortex. Based on *t*-statistic map similarity and *t*-distance RDM similarity the spatial dependencies of this sensitivity are less specifically architectural than when using whitened topographies and crossnobis RDMs. (cf. **Figure 4**). (A) Area p9-46v shows a slightly more positive relationship between geometric and topographic similarity (z-Fisher transformed) than when using whitened contrasts. (B) Transmodal brain areas show a slightly more positive sensitivity of *t*-statistic derived similarities in general, which was absent or reduced with spatially whitened topographies and crossnobis distances. (C) All of the effects identified with whitened metrics are also identifiable with *t*-statistic derived similarity metrics. Associations (right) are corrected for tSNR and test-retest reliability (also estimated using *t*-statistics in this case). *, $p < 0.05$, FDR q = 0.05 jackknife-adjusted spin test. Error bars: 95% confidence interval.

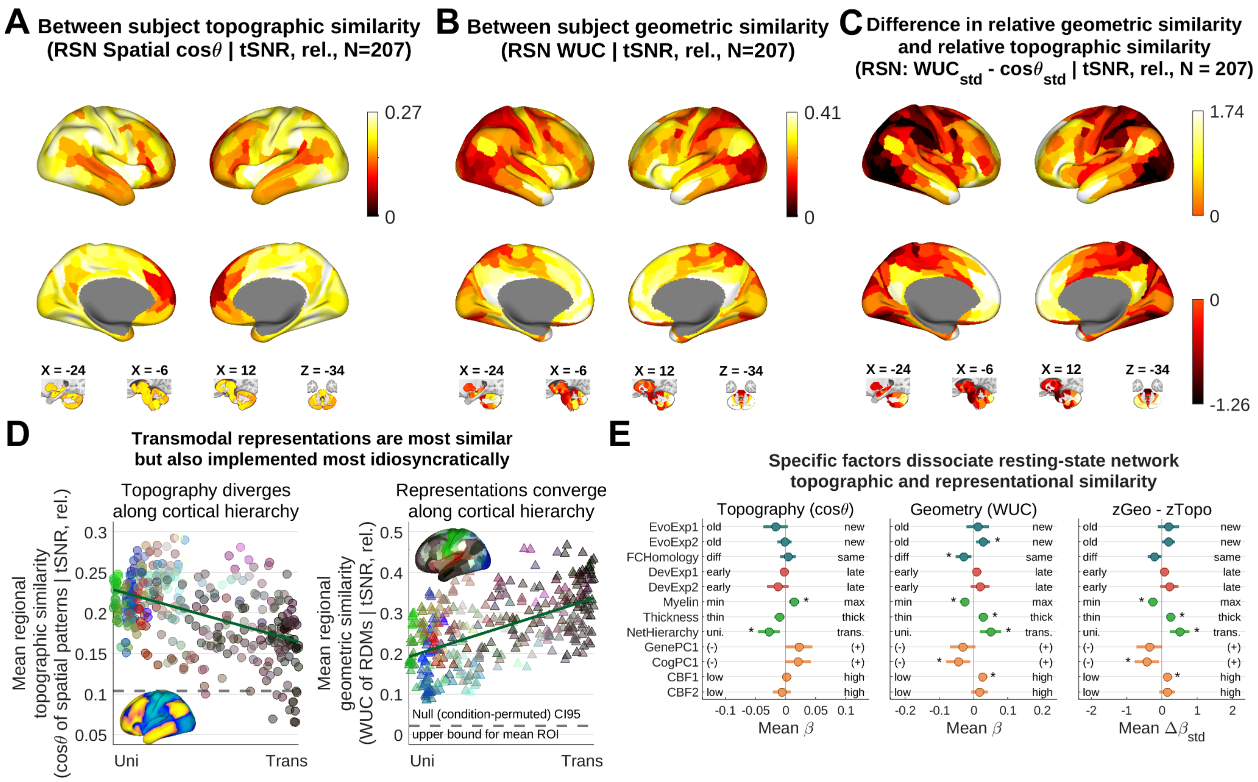


**Supplemental Figure 9** Resting state networks show a nearly identical overall relationship between topography, geometry and neuromaps when using *t*-statistic derived similarity measures as when using whitened topographies and crossnobis RDMs (cf. **Figure 5**). (A) Topographic similarity is uniformly higher among *t*-statistic maps than whitened maps. (B) Geometric similarity of *t*-distance RDMs is indistinguishable from crossnobis RDMs. (C) The difference between topographic and geometric similarity using *t*-statistics is nearly indistinguishable from the same when using whitened topographies and crossnobis RDMs. (D) Qualitatively, *t*-statistic similarity and crossnobis similarity showed a double dissociation with network hierarchy. However, the significant negative association between topographic similarity and network hierarchy is the result of an effect that's only slightly larger than in whitened RSN topographies. (E) RSN *t*-statistic maps show significantly greater topographic similarity in unimodal regions than transmodal regions, in thinner cortex than thicker, and in more myelinated than less myelinated cortex. These effects were not identified with whitened topographies. Geometric similarity and the difference between topography and geometry showed the same relationship with neuromaps when using *t*-statistic derived measures as with whitening (right). Together *t*-statistics based measures show a double dissociation of topographic and representational similarity across multiple gradients. All effects corrected for tSNR and test-retest reliability (estimated based on *t*-statistic similarity in this case). *, $p < 0.05$, FDR q = 0.05 jackknife-adjusted spin test. Error bars: 95% confidence interval.
